# Cobamide sharing drives skin microbiome dynamics

**DOI:** 10.1101/2020.12.02.407395

**Authors:** Mary Hannah Swaney, Shelby Sandstrom, Lindsay R Kalan

## Abstract

The human skin microbiome is a key player in human health, with diverse functions ranging from defense against pathogens to education of the immune system. While recent studies have begun to shed light on the valuable role that skin microorganisms have in maintaining a healthy skin barrier, a detailed understanding of the complex interactions that shape healthy skin microbial communities is limited. Cobamides, the vitamin B_12_ class of cofactor, are essential for organisms across the tree of life. Because this vitamin is only produced by a limited fraction of prokaryotes, cobamide sharing has been shown to mediate community dynamics within microbial communities. Here, we provide the first large-scale unbiased metagenomic assessment of cobamide biosynthesis and utilization in the skin microbiome. We show that while numerous and diverse taxa across the major bacterial phyla on the skin are cobamide dependent, relatively few species encode for *de novo* cobamide biosynthesis. We find that cobamide sharing shapes the network structure in microbial communities across the different microenvironments of the skin and that changes in community structure and microbiome diversity are driven by the abundance of cobamide producers in the *Corynebacterium* genus, in both healthy and disease skin states. Lastly, we find that *de novo* cobamide biosynthesis is enriched only in host-associated *Corynebacterium* species, including those prevalent on human skin. We confirm that the cofactor is produced in excess through quantification of cobamide production by skin-associated species isolated in the laboratory. Taken together, our results support a role for cobamide sharing within skin microbial communities, which we predict stabilizes the microbiome and mediates host interactions.

## INTRODUCTION

The human skin supports a diverse and complex ecosystem of bacterial, fungal, viral, and microeukaryote species, termed the skin microbiome. Highly adapted to live on the skin, these microorganisms form distinct and specialized communities across the skin’s sebaceous, moist, dry, and foot microenvironments. The skin microbiome plays a significant role in human health through contributing to immune system education and homeostasis, protecting against pathogen colonization, and promoting barrier maintenance and repair (Belkaid and Harrison, 2017; Constantinides et al., 2019; Di Domizio et al., 2020; Linehan et al., 2018; Scharschmidt et al., 2017; Wanke et al., 2011).

The transition from taxonomic characterization of the skin microbiome towards study of the mechanisms driving microbe-microbe and microbe-host interactions has shed light on the truly complex nature of skin microbial communities. Recent work has demonstrated that skin commensals not only take part in synergistic and competitive interactions with other microbes (Christensen et al., 2016; Claesen et al., 2020; Nakatsuji et al., 2017; O’Sullivan et al., 2019; Wollenberg et al., 2014), but also participate in host interactions that can dictate skin health and function (Brandwein et al., 2017; Gallo and Nakatsuji, 2011; Naik et al., 2012; Scharschmidt et al., 2017; Uberoi et al., 2021). While these studies have provided fundamental insight into the roles that certain skin commensals, particularly *Staphylococcus* species and *Cutibacterium acnes*, play on the skin, our understanding of the forces that promote stability and mediate overall microbiome structure on healthy skin is still limited.

Within microbial communities, microorganisms interact at a fundamental level through the competition, acquisition, and sharing of nutrients. Nutritional interdependence, for example when one member produces a nutrient that is essential for another, has the potential to impact not only individual species dynamics, but also higher-level interactions, dictating microbial community organization, stability, and function. Of particular interest within microbial communities is sharing of the vitamin B_12_ family of cobalt-containing cofactors, cobamides. Here, we use sharing, as previously defined by Sokolovskaya *et al*., to mean the release of a nutrient or metabolite that is acquired and used by another microorganism (Sokolovskaya et al., 2020).

Cobamides are only synthesized *de novo* by a small fraction of bacteria and archaea, whereas the cofactor is essential for organisms across all domains of life, apart from land plants and fungi. They function in the catalysis of diverse enzymatic reactions, ranging from primary and secondary metabolism, including methionine and natural product biosynthesis, to environmentally impactful processes such as methanogenesis and mercury methylation (Sokolovskaya et al., 2020). Across bacteria, an estimated 86% of bacterial species have been found to encode at least one cobamide-dependent enzyme, whereas only 37% of all bacteria are predicted to produce the cofactor *de novo* (Shelton et al., 2019), suggesting that the majority of bacteria must acquire this important molecule externally. In addition, a unique feature of cobamides is their chemical diversity and functional specificity, with different microorganisms having distinct cobamide preferences and requirements. As such, numerous mechanisms exist for acquisition and use of preferred cobamide(s), including cobamide-specific gene regulation and selectivity by cobamide-dependent enzymes and transporters (Sokolovskaya et al., 2020). Therefore, considering the widespread dependence of cobamides, their limited biosynthesis across bacteria and archaea, and varying specificity organism-to-organism, cobamide sharing is hypothesized to be a major driver of microbial community dynamics. Indeed, *in vitro* and *in vivo* studies of microbial communities, including the human gut microbiome, have demonstrated that cobamide addition modulates community structure, cobamide composition, and expression of cobamide-related genes (Kelly et al., 2019; Men et al., 2015, 2017; Xu et al., 2018; Zhu et al., 2019). In the skin microbiome, however, the role of cobamides has never before been explored.

In the present study, we analyzed 1176 healthy skin metagenomes to predict cobamide dependence and biosynthesis within the skin microbiome and find that phylogenetically diverse skin taxa are predicted to use cobamides, while only a small fraction of species can produce this essential cofactor *de novo*. Modelling of microbial networks shows that cobamide producers, users, and non-users form associations, suggesting a role for cobamide producers that extends beyond direct cobamide sharing. In addition, analysis of taxonomic data from skin metagenomes of four independent studies, including healthy and diseased skin samples, revealed that the abundance of cobamide-producing *Corynebacterium* species is associated with higher microbiome diversity, a key feature of skin health. Lastly, a comparative genomics analysis of 71 *Corynebacterium* species, representing diverse host and environment niche ranges, shows that *de novo* cobamide biosynthesis is almost exclusively present in genomes of host-associated species. Taken together, our results suggest that within the skin microbiome, cobamide sharing is a critical mediator of community dynamics and may play a role in host-microbe interactions through promotion of microbiome diversity

## RESULTS

### Cobamide biosynthesis and precursor salvage genes are encoded by select skin taxa

The *de novo* cobamide biosynthesis pathway is highly complex, consisting of at least 25 enzymatic steps that can be divided into subsections, including tetrapyrrole precursor synthesis, aerobic or anaerobic corrin ring synthesis, nucleotide loop synthesis, and lower ligand synthesis (Figure 1). To determine if cobamide biosynthesis occurs within the skin microbiome, we queried cobamide biosynthesis genes in 1176 skin metagenomes, encompassing samples from 22 distinct skin sites of 3 independent skin microbiome surveys, including the present study, Oh *et al*. (Oh et al., 2016), and Hannigan *et al*. (Hannigan et al., 2015). Using profile HMMs representing 12 genes within the *de novo* cobamide biosynthesis pathway, cbiZ as a marker of cobamide remodeling, and single-copy core gene rpoB as a marker of community structure, we found that samples from sebaceous sites harbored the overall highest median number of hits to cobamide biosynthesis genes, followed by dry, moist, and foot samples (Supplemental Figure 1A).

**Figure 1.**
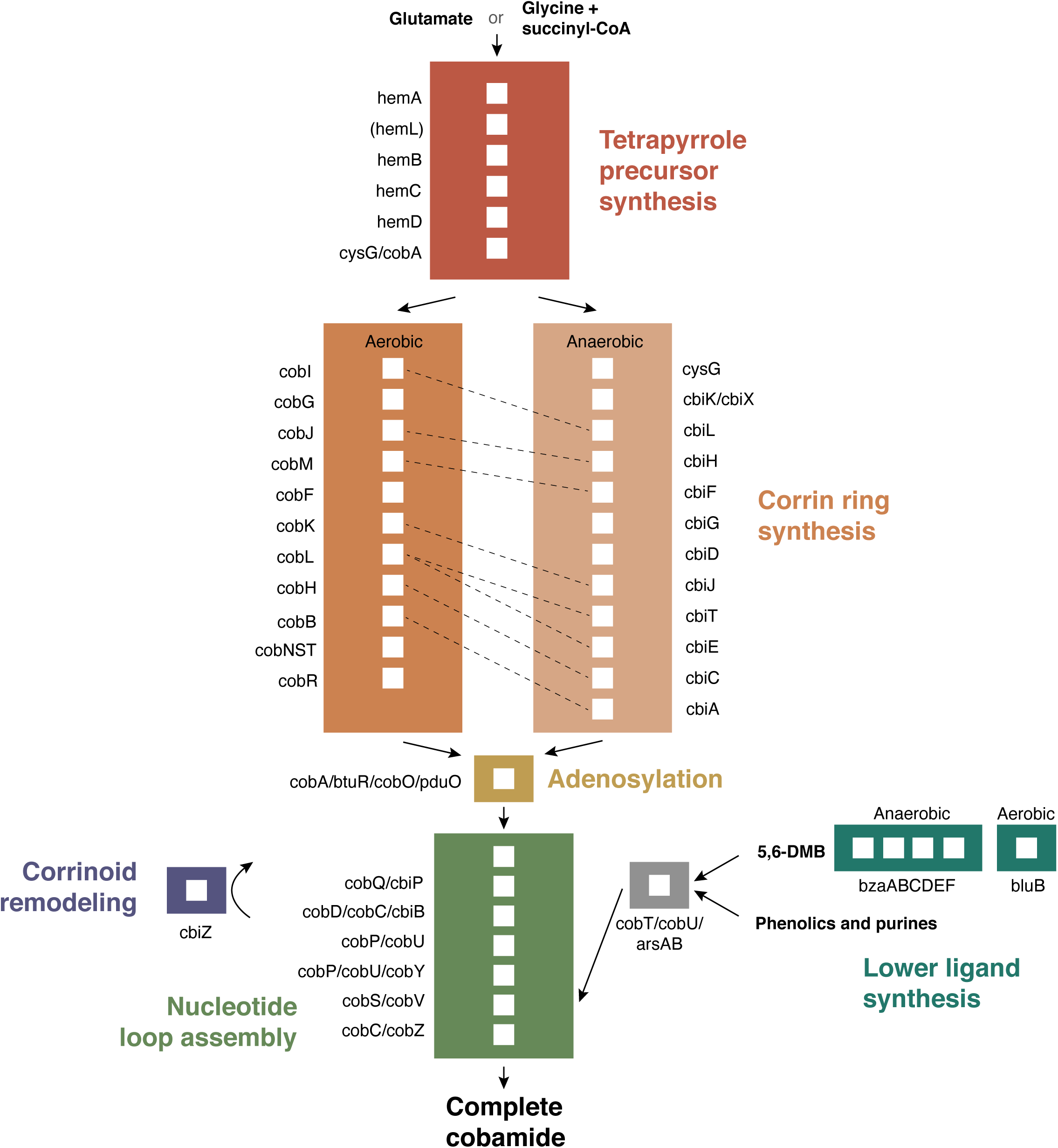
Simplified de novo cobamide biosynthesis pathway. Subsections of the path-way are indicated by color, with gene names and white boxes indicating each enzymatic step. Aerobic and anaerobic corrin ring synthesis pathways contain orthologous enzymes that are indicated with dashed lines. HemL in parentheses is required for synthesis from glutamate.

To assess the contribution of different taxa to cobamide biosynthesis, the metagenomic sequence classifier pipeline Kraken and Bracken was used to classify the resulting gene hits. The top taxa encoding for biosynthetic genes in descending order were determined to be Propionibacteriaceae, Corynebacteriaceae, Veillonellaceae, Streptococcaceae, Dermacoccaceae, and Pseudomonadaceae. Within individual metagenomes, the contribution of each taxon to cobamide biosynthesis gene hits was calculated by dividing the number of biosynthesis gene hits assigned to a given taxa by the total number of biosynthesis gene hits within the sample. We found that Propionibacteriaceae was the dominant contributor to cobamide biosynthesis, particularly in sebaceous sites (Figure 2A).

**Figure 2.**
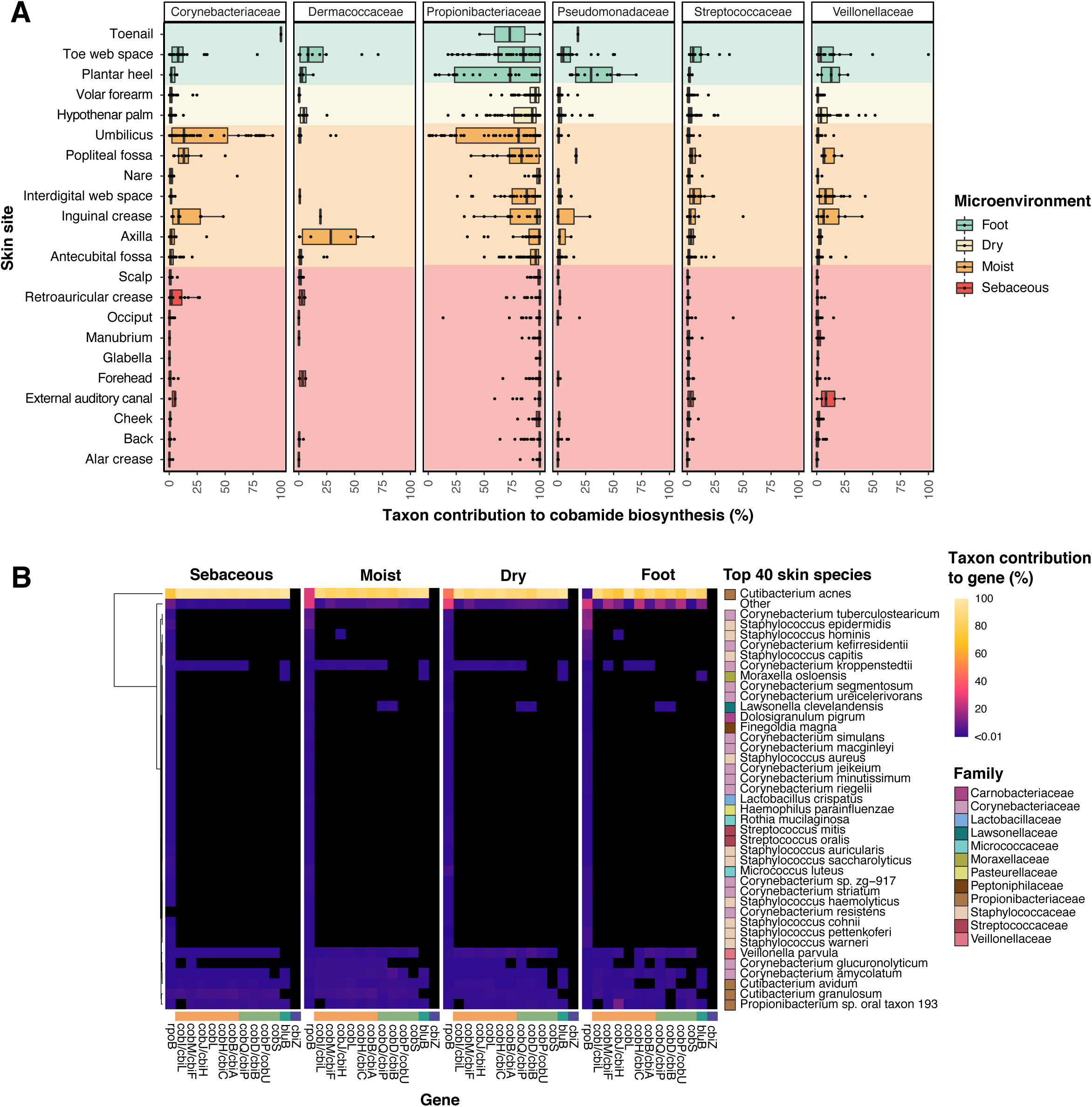
De novo cobamide biosynthesis is limited to distinct taxa on the skin. A) Taxon contribution reflects the proportion of normalized cobamide biosynthesis gene hits assigned to each taxon out of the total normalized cobamide biosynthesis gene hits within a sample. Normalization was performed by dividing hits to each gene by its profile HMM length. Taxon contributions are shown for the top 6 taxa, grouped by skin site. Color indicates microenvironment classification. B) The top 40 most abundant bacterial species within the dataset were determined by totaling the hits to single copy gene rpoB for each species. The remaining species were grouped into “Other”. Individual values in the heatmap represent the number of hits assigned to the species for a particular cobamide biosynthesis gene divided by the total number of hits to the gene. Gene hits were normalized by profile HMM length and sequencing depth prior to calculation. Black squares represent taxonomic abundance from 0 to 0.01%. The colored bar above cobamide biosynthesis genes indicates pathway subsection from Figure 1.

For a finer resolution of taxon contribution, we determined the species contribution of 12 core cobamide biosynthesis genes for the top 40 abundant species within the dataset. Species encoding the full or nearly complete suite of cobamide biosynthesis markers was consistent across microenvironments, with *Cutibacterium acnes* being the dominant contributing species (Figure 2B). Other species contributing to most or all of the biosynthesis markers include *Corynebacterium amycolatum, Corynebacterium kroppenstedtii, Corynebacterium glucuronolyticum, Veillonella parvula, Cutibacterium granulosum*, and *Propionibacterium* sp. oral taxon 193. A proportion of low abundance skin taxa were predicted to encode for the full suite of cobamide biosynthesis markers (grouped into the “Other” category), demonstrating that cobamide biosynthesis is encoded by both dominant and rare taxa.

Taxa such as Moraxellaceae and Xanthomonadaceae encode for a limited set of cobamide biosynthesis genes including cobQ/cbiP, cobD/cbiB, cobP/cobU and cobS (Supplemental Figure 2), which can function in cobamide precursor salvage (Gray et al., 2008; Rodionov et al., 2019). This suggests that cobamide synthesis through salvaging is also occurring on the skin. Furthermore, cobamides are grouped into three classes based on their structurally distinct lower ligand: benzimidazolyl, purinyl, and phenolyl cobamides (Sokolovskaya et al., 2020). Most predicted cobamide producers identified in this analysis likely synthesize benzimidazolyl cobamides because they encode for bluB, the gene responsible for the aerobic synthesis of lower ligand 5,6-dimethylbenzimidazole (DMB) (Figure 2B, Supplemental Figure 2) (Campbell et al., 2006; Gray and Escalante-Semerena, 2007; Taga et al., 2007). However, in species such as *V. parvula, C. granulosum*, and *C. glucuronolyticum*, bluB is absent, suggesting that these species produce non-benzimidazolyl cobamides. Overall, our results demonstrate that select taxa within the skin microbiome have the genetic potential to produce chemically diverse cobamides through *de novo* biosynthesis or precursor salvage and that *de novo* biosynthesis is restricted to only a few species.

### Phylogenetically diverse skin taxa are cobamide dependent

Although few species within the skin microbiome synthesize cobamides *de novo*, we predict that a larger proportion use cobamides. We determined the prevalence of the cobamide transport protein btuB and 19 enzymes that carry out diverse cobamide dependent reactions. The median number of cobamide-dependent gene hits across samples varied by microenvironment (Supplemental Figure 1). Across the sebaceous, moist, and dry microenvironments, Propionibacteriaceae was the dominant family encoding for the cobamide-dependent enzymes D-ornithine aminomutase, methylmalonyl-CoA mutase, and ribonucleotide reductase class II (Figure 3). In contrast, across the remaining cobamide dependent enzymes, hits were assigned to phylogenetically diverse taxa across the four major phyla on the skin (Actinobacteria, Firmicutes, Proteobacteria, and Bacteroidetes) (Grice and Segre, 2011). Cobamide-dependent enzymes involved in primary metabolism, including methionine synthase, epoxyqueosine reductase, ribonucleotide reductase, and ethanolamine lyase, were the most common cobamide dependent enzymes in the dataset (Supplemental Figure 3). Notably, only 1% of species appreciably contribute to de novo cobamide biosynthesis (n=18 species), yet approximately 39% of species encode for cobamide dependent enzymes (n=638 species encoding at least one cobamide-dependent enzyme) (Supplemental Figure 3). While the true number of *de novo* cobamide producers may be underestimated due to filtering of rare and singleton hits prior to analysis, these species likely represent the core cobamide producers found on the skin. Overall, these results support a model of cobamide sharing, where a much larger number of skin taxa require cobamides than can produce the cofactor *de novo*.

**Figure 3.**
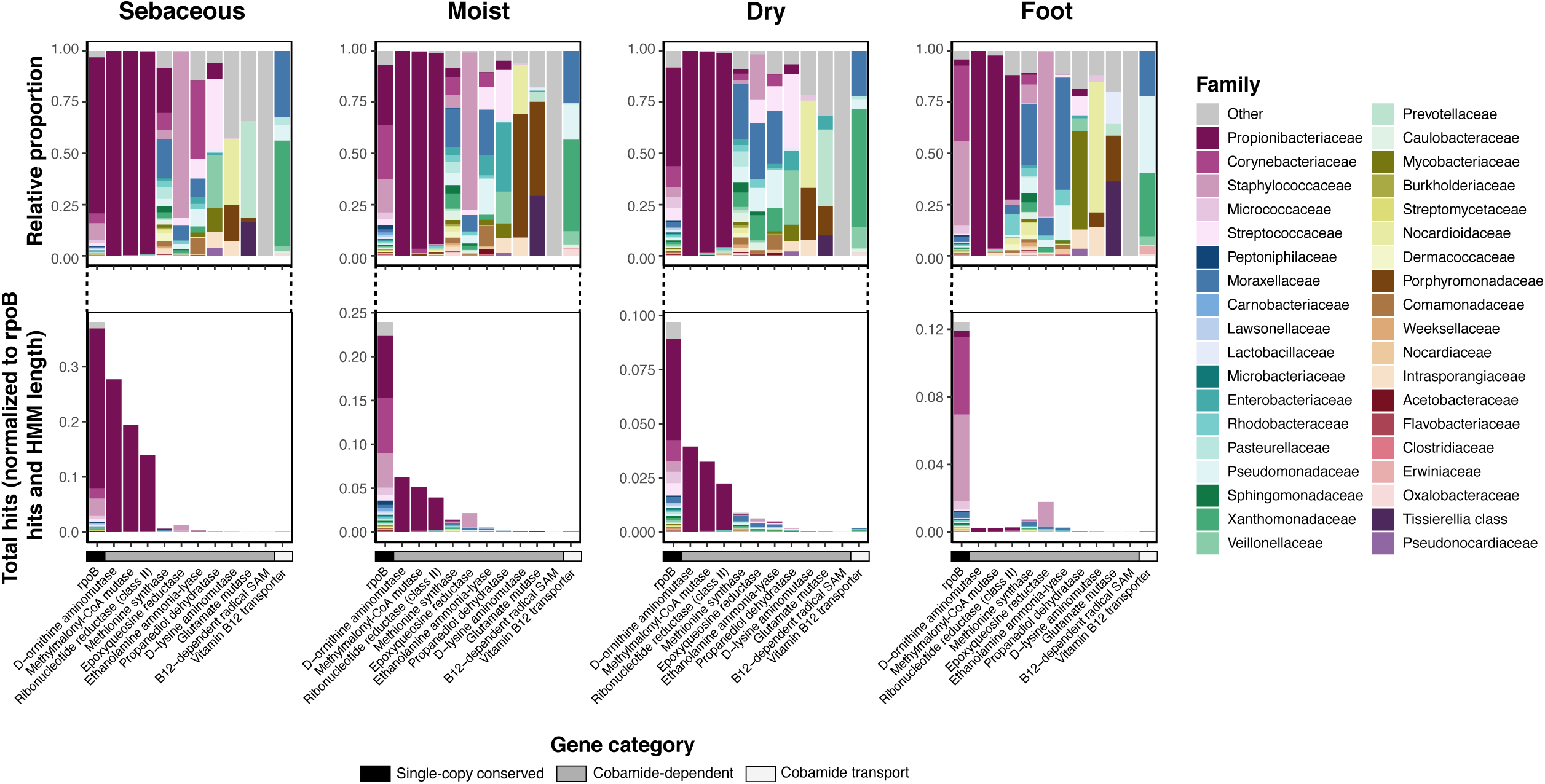
Phylogenetically diverse skin bacteria encode for cobamide dependent enzymes and transporters. The total normalized hits for cobamide-dependent enzymes, cobamide transport protein btuB, and SCG rpoB are shown (total hits normalized to profile HMM coverage and sequence depth), with the taxonomic abundance of the hits expanded as relative proportions above. Hits to distinct B12-dependent radical SAM proteins are grouped together as “B12-dep radical SAM”.

### Regulation of cobamide biosynthesis is species-specific

To further delineate cobamide usage within the skin microbiome, we identified cobalamin riboswitches within the metagenomes. Cobalamin riboswitches are cobamide-binding elements found in the untranslated region of bacterial mRNAs that regulate expression of genes or transcripts involved in cobamide-dependent metabolism, biosynthesis, and cobamide transport (Garst et al., 2011; Nahvi et al., 2004; Polaski et al., 2017). We show that phylogenetically diverse skin taxa encode for cobalamin riboswitches, with Propionibacteriaceae being the dominant taxa (Figure 4A). At the species level, these hits were found predominantly within *C. acnes* genomes.

**Figure 4.**
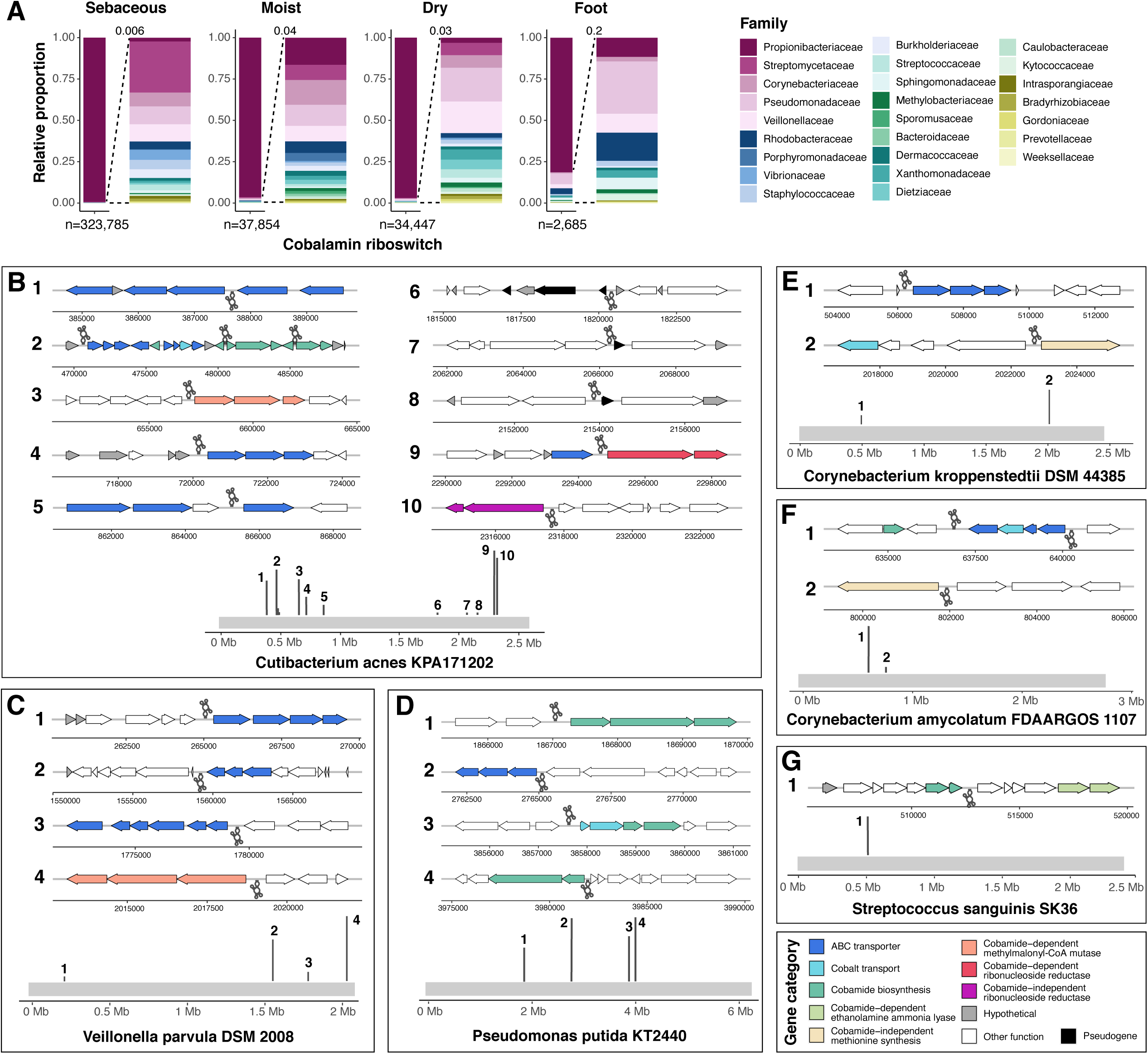
Cobalamin riboswitch regulation varies across skin taxa. A) The taxonomic abundance of hits for cobalamin riboswitches (Rfam clan CL00101) are shown, with an expanded view of low abundance hits to the right. Total cobalamin riboswitch hits within each microenvironment are indicated. Cobalamin riboswitch-containing reads identified from INFERNAL analysis were aligned to B) Cutibacterium acnes KPA171202, C) Veillonella parvula DSM 2008, D) Pseudomonas putida KT2440, E) Corynebacterium kroppenstedtii DSM 44385, F) Corynebacterium amycolatum FDAARGOS 1107, and G) Streptococcus sanguinis SK36 genomes. Dark gray lines along the light grey genome track indicate the position of mapped INFERNAL hits within the genome. Genes upstream and downstream of the riboswitches are colored by their general functional annotation. White (other function) indicates genes not currently known to be associated with cobamides. Grey (hypothetical) indicates a hypothetical protein that has no functional annotation. Right-facing gene arrows and upright dark gray riboswitch icons indicate for-ward strand orientation, and left-facing gene arrows and inverted riboswitch icons indicate reverse strand orientation. Genomic regions are not to scale.

To identify the pathways regulated by cobamides in *C. acnes*, we mapped cobalamin riboswitch sequence reads to the *C. acnes* KPA171202 reference genome. We find that riboswitches are distributed across the genome in numerous regions, regulating pathways involved in ABC transport, cobalt transport, cobamide biosynthesis, and cobamide-dependent and -independent reactions (Figure 4B). Three of the *C. acnes* cobalamin riboswitches (Regions 6, 7, and 8) are located upstream of pseudogenes or genes of unknown function (Figure 4B). Manual curation of these sequences suggest that the small pseudogenes are hypothetical adhesin protein fragments and the larger downstream sequences are thrombospondin type-3 repeat containing proteins (Supplemental Material S4). The role of cobalamin riboswitches in regulation of these genes is unknown, but suggests that cobalamin riboswitches are regulating diverse functions yet to be discovered.

We found fewer cobalamin riboswitches in the genomes of other species relative to *C. acnes*. Riboswitches were identified near gene neighborhoods with functions involved in cobamide biosynthesis, ABC transport, cobalt transport, and both cobamide-dependent and cobamide-independent isozymes (Figure 4C-G). Unlike *C. acnes*, which tightly regulates cobamide biosynthesis, our data do not support riboswitch regulation of cobamide biosynthesis in *V. parvula, C. kroppenstedtii*, and *C. amycolatum*, suggesting constitutive *de novo* production of the molecule occurs on the skin. Overall, cobalamin riboswitches are likely to regulate diverse processes in the skin microbiome, including novel functions.

### Cobamide biosynthesis and usage shapes microbial network structures

Having determined that cobamide producers, precursor salvagers, and users are prevalent within the skin microbiome, we sought out to determine how these members may be interacting, both with each other and with members who neither use nor produce cobamides. We utilized the SPIEC-EASI statistical method to infer microbial associations between common species on the skin (see Supplemental Material S5). Associations present in at least two of the three metagenomic studies were used to generate a final consensus network for each skin microenvironment.

Across all microenvironments, the majority of associations are positive, with few negative associations in each network (Figure 5A). The following measurements were used to quantify each network: node degree, density, transitivity, modularity, and phylum assortativity (see Supplemental Table 1 for a description of these properties). Overall, the moist environment network was the least sparse and modular and the most dense and transitive, suggestive of a more interconnected and less modular community. The dry network was the most sparse and modular and the least dense and transitive, suggesting the existence of interaction modules with dense connections between species of the same module. Sebaceous and foot networks fell in the middle of this spectrum. Across all microenvironments, we observed high assortativity by phylum, indicating a preference for species to associate with other species in the same phylum.

**Figure 5.**
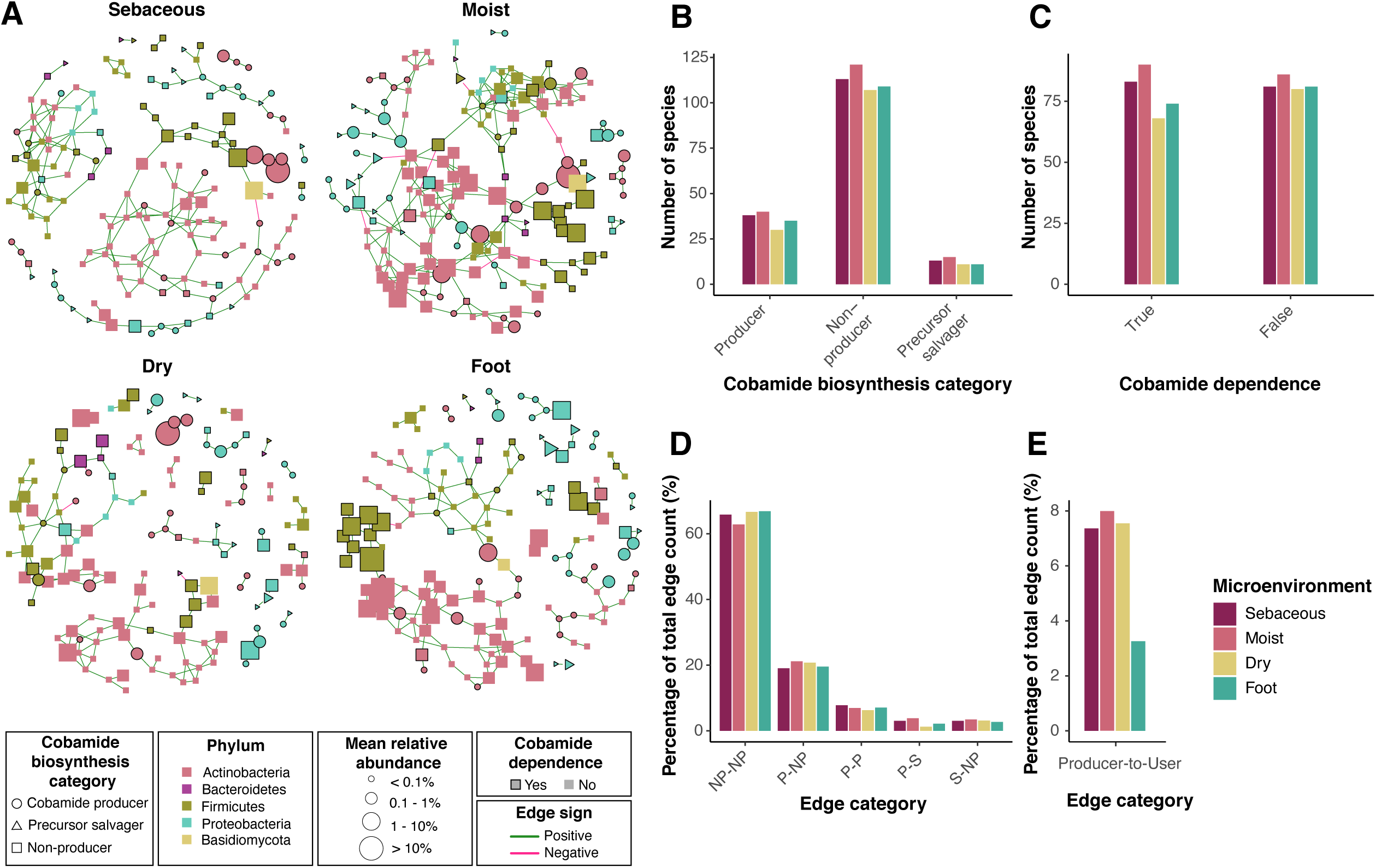
Skin microbiome networks reveal microbial associations among cobamide producers, precursor salvagers, and users. A) The SPIEC-EASI method was used to identify microbial associations within each microenvironment of three independent skin microbiome datasets. Consensus networks are shown, representing associations identified in at least 2 of the 3 datasets. Species are represented by nodes and colored by phylum. Green and pink edges represent positive and negative associations, respectively. Node shape represents cobamide biosynthesis category and node size reflects mean species relative abundance within each microenvironment. Cobamide dependent species are outlined in black. In each final network, B) the number of species classified to each cobamide biosynthesis category, C) the number of species that are cobamide dependent or independent, D) the percentage of total edges that fall into each cobamide biosynthesis edge category, and E) the percentage of total edges that exist between cobamide producers and cobamide dependent species that are non-producers or precursor salvagers is shown. NP=Non-producer, P=Producer, S=Precursor salvager.

Across microenvironments, the distribution of species identified to be cobamide producers, precursor salvagers, and non-producers was relatively consistent, with more non-producers than producers or precursor salvagers (Figure 5B). The distribution of species identified to be cobamide dependent or not cobamide dependent was also consistent across microenvironments, with a generally equal number of cobamide-dependent species to cobamide-independent species in each network (Figure 5C). Edges were quantified based on cobamide biosynthesis category, showing more non-producer to non-producer edges, followed by producer to non-producer, producer to producer, producer to precursor salvager, and lastly precursor salvager to non-producer (Figure 5D). The moist microenvironment has the largest number of edges between species that are *de novo* producers and those that are non-producers yet cobamide-dependent, followed by sebaceous, dry, and foot microenvironments. Overall, we find that associations between cobamide producers, precursor salvagers, non-producers, users, and non-users are distributed throughout the networks, suggesting that cobamide sharing between users and producers can impact microbial interactions within the entire network.

### Microbial diversity and community structure is driven by cobamide producer abundance

While the majority of bacteria are predicted to encode at least one cobamide-dependent enzyme, only 37% of bacteria are predicted to produce the cofactor *de novo (Shelton et al*., *2019)*. Therefore within microbial communities, cobamide sharing likely exists as a means to fulfill this nutritional requirement and is hypothesized to mediate community dynamics. On the skin, two of the top most abundant genera are *Cutibacterium* and *Corynebacterium*, both of which we found to include species that are *de novo* cobamide producers. Therefore, we hypothesize that changes at the community level are associated with the presence of these cobamide-producing species. To assess this, we first explored the relationship between microbiome diversity and cobamide-producing Corynebacteria (CPC) abundance within healthy skin metagenomes. NMDS ordination of Bray-Curtis dissimilarity indices revealed clustering that follows increasing gradients of both alpha diversity and CPC abundance, where alpha diversity increases as CPC abundance increases (Figure 6A, Supplemental Table 2). This was most striking for samples from sebaceous, moist, and foot sites. In contrast, this pattern of clustering was not observed for *Cutibacterium* cobamide producers, but rather samples with the highest *Cutibacterium* relative abundances often were the least diverse (Supplemental Figure 5). Furthermore, communities with a low CPC abundance were usually dominated by *Cutibacterium acnes*, whereas communities with high abundance showed an expansion of other skin taxa and an overall more even species distribution within the community (Supplemental Figure 6). Consistent with our analysis of riboswitch regulation, these results further support a model where *Corynebacterium* species constitutively produce cobamides as a shared common good, promoting microbiome diversity and structure. On the other hand, tightly regulated production by *Cutibacterium* species permits niche expansion and lower diversity.

**Figure 6.**
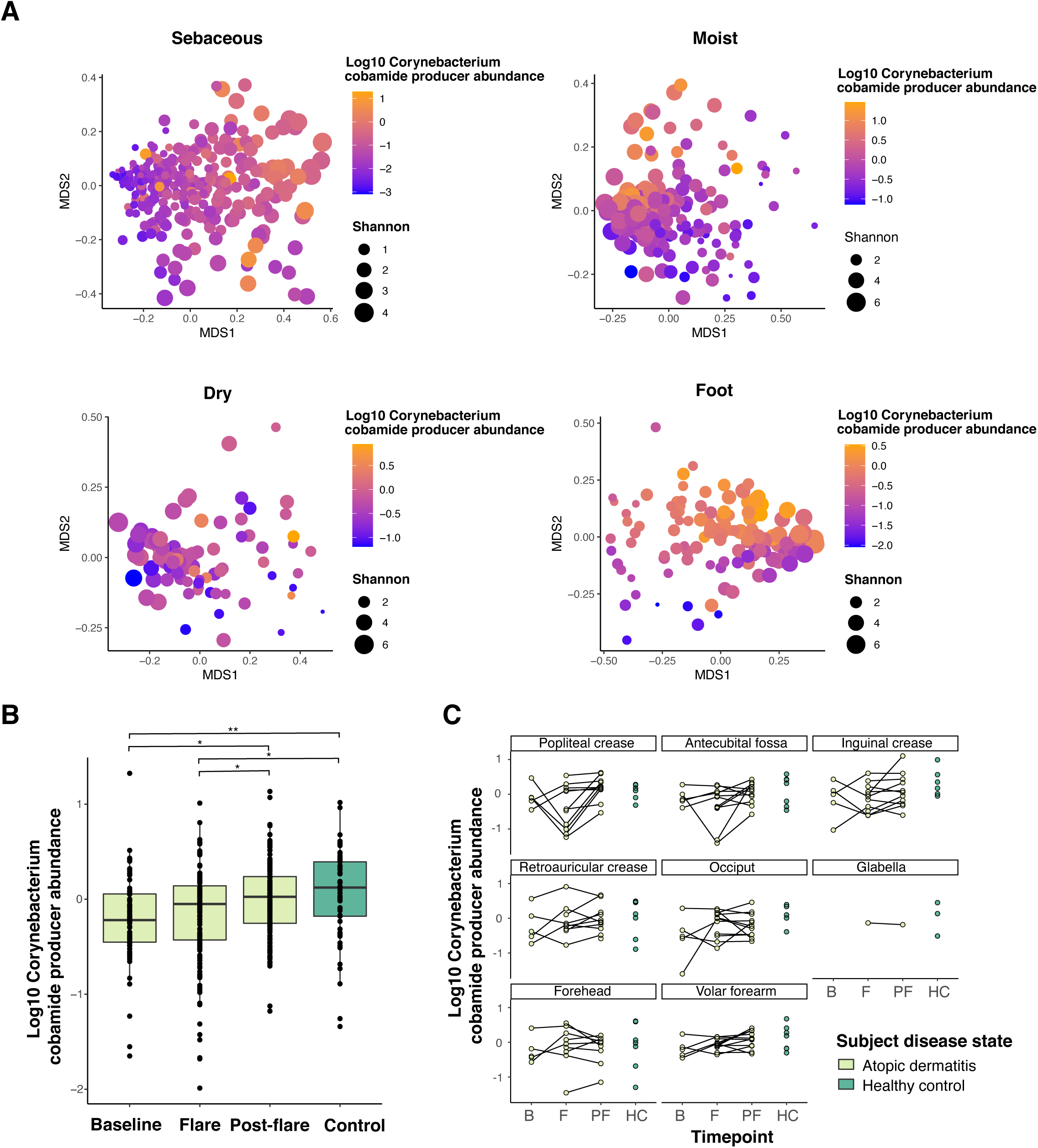
Cobamide-producing Corynebacterium abundance is associated with microbiome diversity and atopic dermatitis disease state. Within each metagenome, the cumulative relative abundance of cobamide-producing Corynebacteria (CPC) was calculated. A) NMDS plots based on Bray-Curtis indices for healthy adult samples within each skin microenvironment are shown. Points are colored by Corynebacterium cobamide producer relative abundance and sized by alpha diversity (Shannon). B) The relative abundance of CPC in pediatric atopic dermatitis patients at baseline, flare, and post-flare timepoints or in healthy control subjects. A pairwise Wilcoxon rank sum test was performed among each group with FDR correction (*<0.05, **<0.01) (C) The relative abundance of CPC in each individual skin site sampled. Black lines connect time-points for a given patient. Certain sites were sampled from both sides of the body, therefore each point represents the average abundance of for each individual at the specified skin site.

### Cobamide production is depleted in atopic dermatitis

A decrease in microbiome diversity is associated with increased pathogen colonization in dermatological disease such as atopic dermatitis (AD) (Paller et al., 2019; Williams, 2005). To assess the potential role of CPC in the AD skin microbiome, we analyzed 417 metagenomes from a cohort of 11 pediatric AD patients and 7 healthy controls. Microbiome structures exhibited a higher level of variability compared to the adult cohorts, with weak clustering of samples based on alpha diversity or CPC abundance. A subset of samples collected from moist sites during a flare formed a distinct cluster exhibiting low CPC abundance and alpha diversity (Supplemental Figure 7). AD skin symptoms often present in moist sites such as the antecubital fossa (bend of the elbow) and popliteal fossa (bend of the knee), suggesting a relationship between microbiome structure, diversity, and CPC abundance during AD flares. Consistent with this hypothesis, we observed that CPC abundance is significantly reduced in AD patients at baseline (p=0.0018) as well as during flares (p=0.0050) compared to healthy controls (Figure 6B). Within individual patients, a sharp decrease in CPC abundance between baseline and flare occurs in a subset of patients, particularly in the antecubital fossa and popliteal fossa (Figure 6C). Overall, differential CPC abundance is detected between disease states, suggesting a relationship between these members and microbial community structure in atopic dermatitis.

### Cobamide biosynthesis is enriched in host-associated *Corynebacterium* species

Until recently, species of the *Corynebacterium* genus have been underappreciated as significant members of skin microbial communities, predominantly due to the difficulty of growing these species in the lab, which is a result of their nutritionally fastidious and slow-growing nature (Grice and Segre, 2011). However, sequencing efforts have revealed that Corynebacteria are a dominant taxon within the microbiome, particularly in moist skin microenvironments (Grice et al., 2009; Oh et al., 2014, 2016). Our results suggest an important role for cobamide production by skin-associated *Corynebacterium* species. Because other species within the *Corynebacterium* genus occupy highly diverse habitats, including soil, cheese rinds, coral mucus, and other human and animal body sites (Bernard, 2012), we were interested in exploring the genomic diversity within the *Corynebacterium* genus and how it relates to cobamide biosynthesis. To do so, we performed a pangenome analysis using anvi’o, which included 50 host-associated and 21 environment-associated *Corynebacterium* genomes (Supplemental Material S8), acquired as complete assemblies from NCBI (n=68) or as draft assemblies from human skin isolates (n=3). Gene clusters (GCs), which are computed and used by anvi’o, represent one or more genes grouped together based on homology at the translated DNA sequence level (Delmont and Murat Eren, 2018). Across all species, 42,154 total GCs were identified. 495 of these are core GCs present in all genomes, 13,235 GCs are shared (dispensable), and 28,424 GCs are found in only one genome (species-specific) (Supplemental Figures 8-9). Genome size ranged from 2.0 to 3.6 Mbp, with an average of 2.7 ± 0.3 Mbp, and the number of GCs per genome ranged from 1858 to 3170 GCs, with an average of 2365 ± 294 GCs (Supplemental Material S8). Host-associated species have significantly fewer GCs per genome compared to environment-associated species (2174 vs. 2664, p-value<0.0001) and a significantly reduced median genome length (2.52 Mbp vs 3.03 Mbp, p-value<0.0001) (Figure 7B, 7C).

**Figure 7.**
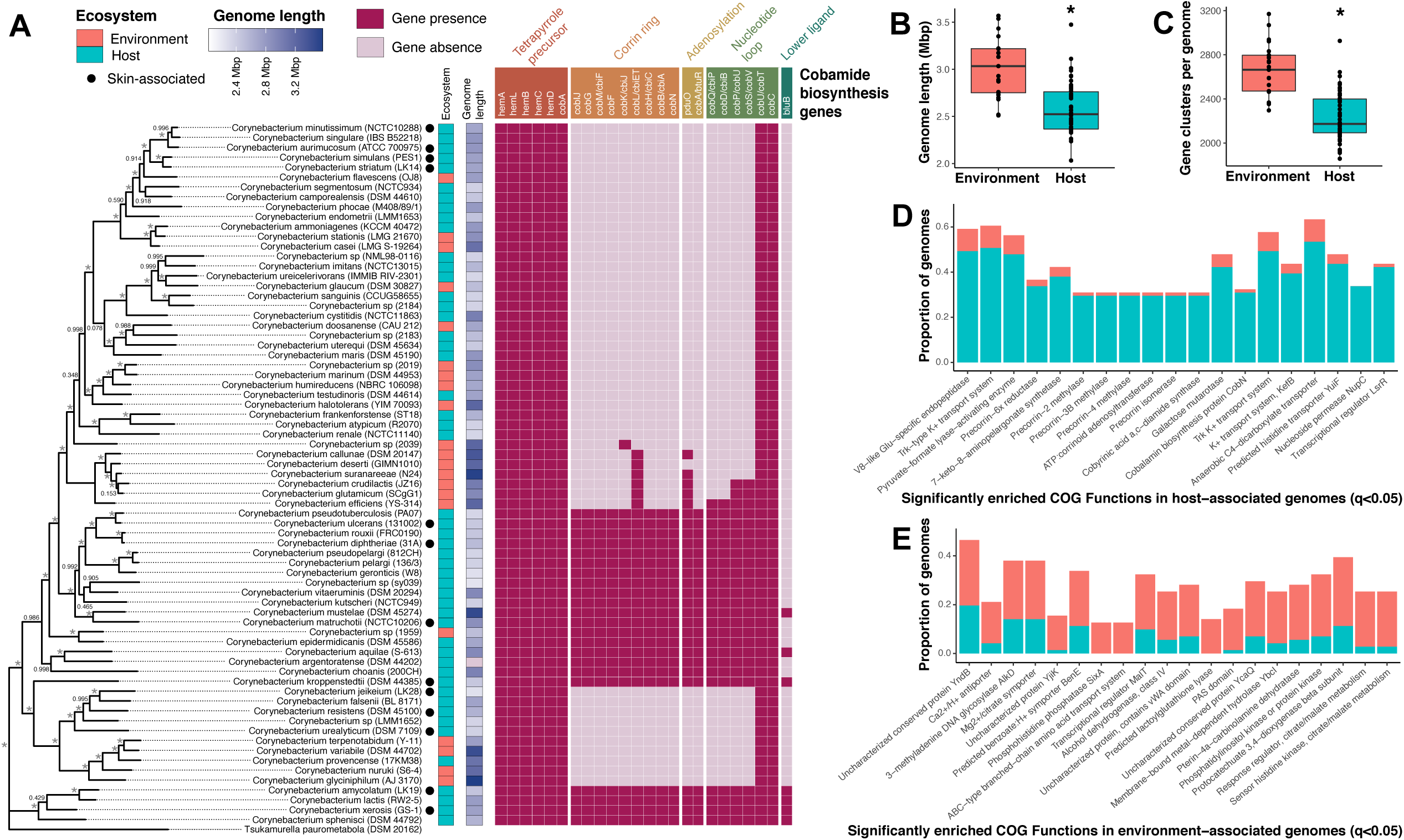
De novo cobamide biosynthesis is host-associated within the Corynebacterium genus. A) A Corynebacterium phylogenetic tree based on comparison of 71 conserved single copy genes was generated using FastTree within the anvi’o environment. The tree is rooted with Tsukamurella paurometabola, and bootstrapping values are indicated (* = 100% bootstrap support). Species are colored by host (blue) or environment (orange) association, and by genome length (dark blue). KOfamScan was used to identify the presence (dark pink) or absence (light pink) of cobamide biosynthesis genes within each genome. Cobamide biosynthesis subsections are indicated and differentially colored based on B) Genome length and C) number of gene clusters for the Corynebacterium genomes were determined using anvi’o. Significantly enriched COG functions in D) host-associated or E) environment-associated genomes were identified with anvi’o. The top 20 significantly enriched COG functions (q < 0.05) are shown, ordered by ascending significance. Blue = host-associated, orange = environment-associated.

We determined functions that differ between host- and environment-associated genomes using a functional enrichment analysis. The top significantly enriched functions in environment-associated genomes include pathways putatively involved in amino acid transport, metabolism of various substrates including aromatic compounds, tetrahydropterin cofactors, and citrate/malate, and other uncharacterized functions (q < 0.05) (Figure 7E). Within host-associated genomes, we observed a significant enrichment of pathways involved in the transport of various metabolites and ions, as well as 8 COG functions involved in cobamide biosynthesis (q < 0.05) (Figure 7D). To identify and validate the presence of the *de novo* biosynthesis pathway within the 71 *Corynebacterium* genomes, we scanned the genomes using KOfamScan. Tetrapyrrole precursor synthesis, which is shared among the cobamide, heme, and chlorophyll biosynthesis pathways (Shelton et al., 2019), was conserved throughout the genus (Figure 7A). Corrin ring and nucleotide loop synthesis was intact and conserved within 5 distinct *Corynebacterium* lineages, including those of *C. diphtheriae, C. epidermidicans, C. argentoratense, C. kroppenstedtii*, and *C. amycolatum*. The species within these groups encode for all or nearly all of the genes required for *de novo* cobamide biosynthesis, and notably, 21 out of 22 of these predicted cobamide producers are host-associated. Taken together, these results demonstrate a range of cobamide biosynthetic capabilities by Corynebacteria, with *de novo* producing species being almost exclusively host-associated, despite reduced genome size. Thus, we hypothesize a role for cobamides in mediating host-microbe interactions.

### Skin commensal *Corynebacterium amycolatum* produces high levels of cobamides

From our metagenomic and comparative genomic analyses, we identified *C. amycolatum* as a *de novo* cobamide producer. To test *in vitro* production of cobamides by this species, we isolated a strain of *C. amycolatum* from healthy skin, cultured it in a minimal growth medium, and prepared cell extracts from the intracellular metabolite content. We tested the cell extract in a microbiological assay using the indicator strain *E. coli* ATCC 14169, whose growth is proportional to cobamide concentration from 0.1 to 1.5 ng/mL (Supplemental Figure 10A). When diluted 10,000- to 50,000-fold, *C. amycolatum* cell extracts yielded growth of *E. coli* within the linear range (Supplemental Figure 10B), with an average cobamide amount of 1.51 ± 0.135 µg per gram of wet cell weight and an average intracellular concentration of 11.3 ± 2.37 µM. Physiological requirements of cobamides range from nanomolar to even picomolar concentrations (Sokolovskaya et al., 2020), supporting our hypothesis that *C. amycolatum* produces the cofactor in excess quantities to support cobamide sharing in the community.

## DISCUSSION

This study, which provides the first in depth analysis of cobamide biosynthesis and use within skin microbial communities, contributes to the growing body of evidence that nutrient sharing is a critical driver of microbial community dynamics. Our analysis of skin metagenomic data demonstrates that phylogenetically diverse skin taxa, both high and low abundance, encode for metabolically diverse cobamide-dependent enzymes, as well as proteins involved in cobamide transport and salvage. Meanwhile, *de novo* producing skin species are greatly outnumbered by the total number of species that require cobamides for metabolism. In contrast to studies of cobamides in other microbial communities that have demonstrated *de novo* synthesis to be carried out by a relatively small fraction of the community (Lu et al., 2020; Magnúsdóttir et al., 2015; Romine et al., 2017), our results indicate that cobamides are produced by taxa considered to be dominant members within the skin microbiome, including *Cutibacterium* and *Corynebacterium* species. However, what is currently unknown is the extent to which cobamides produced by these dominant taxa are available for community use.

Our findings show that regulation of biosynthesis and use can vary drastically from taxa to taxa. For example, we show that *C. acnes* is the top species encoding for cobamide biosynthesis and dependent genes within the dataset, yet expression of these genes is under tight regulation by over ten cobalamin riboswitches. Conversely, other predicted cobamide producers on the skin, such as *C. amycolatum* and *C. kroppenstedtii*, only possess a few cobalamin riboswitches, and these riboswitches regulate cobamide-dependent, -independent, and transport functions, as opposed to cobamide biosynthesis. The absence of riboswitch-regulated biosynthesis genes is similarly observed across all Corynebacteriaceae (Sun et al., 2013). This suggests that constitutive expression of cobamide biosynthesis occurs in specific skin taxa. Overall, cobamide production and riboswitch regulation are likely to act as important mediators of microbe-microbe interactions on the skin.

Within microbial communities, cobamides are hypothesized to mediate community dynamics because of the relative paucity of cobamide producers, yet evident requirement for this cofactor across the bacterial domain of life (Degnan Patrick H. Taga Michiko E. Goodman, 2014; Shelton et al., 2019; Sokolovskaya et al., 2020). Our results suggest that on the skin, *Corynebacterium* cobamide-producing species promote microbiome diversity and dictate community structure in both healthy and diseased skin states. In addition, microbial association analysis identified associations between cobamide producers, users, as well as non-users, revealing the opportunity for cobamide sharing to impact microbiome dynamics at a community level. Because there exists a spectrum of ecological niches on the skin, we propose that in addition to the existence of cobamide sharing, cobamide-mediated interactions are dependent upon the spatial structure of skin microbial communities. For example, *C. acnes* is an anaerobe that predominantly resides deep within the anaerobic sebaceous follicle (Dréno et al., 2018), dominating between 60-90% of the follicle community (Hall et al., 2018). As such, the opportunity for cobamide-mediated interactions is likely reduced as a result of the *C. acnes*-dominated sebaceous gland. Approaching the more oxygenated skin surface, the community becomes more diverse (Oh et al., 2014), thus increasing the incidence of cobamide interactions and subsequent effects on community dynamics.

Corynebacteria are well-equipped for growth on the skin due to their “lipid-loving” and halotolerant nature, allowing them to thrive in moist and sebaceous skin microenvironments (Scharschmidt and Fischbach, 2013). However, many questions remain about the processes that govern skin colonization by this relatively understudied skin taxa and how these processes may impact or be impacted by microbe-microbe and microbe-host interactions on the skin. We identified several *Corynebacterium* species to be *de novo* cobamide producers on the skin, and further, that the abundance of these species impacts microbial community dynamics through promotion of diversity. This suggests skin-associated Corynebacteria are a keystone species, leading us to perform a comparative genomics study of the entire genus. As expected, host-associated species have significantly smaller genomes, but unexpectedly, they are enriched for *de novo* cobamide biosynthesis as compared to environment-associated species. Retention of the energetically costly 25-enzyme cobamide biosynthesis pathway within host-associated species, even with reduced genome size, suggests that synthesis of this cofactor is advantageous for host niche colonization.

A key question that arises is why some *Corynebacterium* species have retained the *de novo* cobamide biosynthesis pathway, while others have not. Our results showed that *Corynebacterium* species encode for cobamide-dependent methionine synthase, methylmalonyl-CoA mutase, and ethanolamine ammonia lyase, consistent with previous findings by Shelton *et al*. (Shelton et al., 2019). Therefore, cobamides are likely produced by Corynebacteria to fulfill metabolic requirements in methionine, propionate, and glycerophospholipid metabolism. Alternative cobamide-independent pathways exist for these functions, therefore cobamides may confer a distinct advantage for these species. Indeed, metE, the cobamide-independent methionine synthase, is sensitive to oxidative stress and has reduced turnover compared to metH (González et al., 1992; Hondorp and Matthews, 2004; Leichert and Jakob, 2004). The skin in particular is subject to high oxidative stress as a result of metabolic reactions, cosmetics, and UV irradiation exposure. (Andersson et al., 2019; Hakozaki et al., 2008; Kawashima et al., 2018). Therefore, while the significance of employing cobamide-dependent vs -independent isozymes for bacterial metabolism on the skin is unknown, inherent features of the skin such as high oxidative stress may play a role.

Our study further elucidates a potentially novel cobamide-mediated host-microbe interaction. We determined that 20 of 21 *Corynebacterium* species encoding for *de novo* biosynthesis are host-associated. Most of these species are characterized by their colonization of epithelial-associated sites (skin, oral cavity, nasal cavity) of their various hosts, including humans, other small and large mammals, and birds. While a clear role for cobamides in host-microbe interactions is not currently defined, Kang *et al*. have demonstrated that in humans, oral vitamin B_12_ supplementation can repress cobamide biosynthesis genes of the skin microbiota, thus providing evidence that host-acquired cobamides are available to the microbiota through the skin (Kang et al., 2015). Whether microbially-produced cobamides are accessible to the host through these epithelial surfaces is unknown, but warrants future investigation.

Our findings reveal that cobamide dependence is widespread across the phylogenetic diversity of the skin microbiome, while a small number of skin taxa are capable of *de novo* production, including several species of the *Corynebacterium* genus. Within skin microbial communities, abundance of these cobamide-producing Corynebacteria is strongly associated with increased microbiome diversity and disease state, supporting our hypothesis that cobamides are important mediators of microbiome structure and skin health. We also show that within the *Corynebacterium* genus, *de novo* cobamide biosynthesis is uniquely a host-associated function. Future studies to interrogate the role of cobamides in microbe-microbe and microbe-host interactions will provide insight into the key roles that microbially-derived metabolites play in microbial community dynamics and host health.

## Acknowledgements

This work was supported by grants from the National Institutes of Health (NIAID U19AI142720 [L.R.K], NIAID T32AI055397 [M.H.S]). The content is solely the responsibility of the authors and does not necessarily represent the official views of the National Institutes of Health. This work used the Extreme Science and Engineering Discovery Environment (XSEDE), which is supported by National Science Foundation grant number ACI-1548562. Specifically, it used the Bridges system at the Pittsburgh Supercomputing Center (PSC) through allocation ID MCB200172. The authors gratefully acknowledge Dr. Michi Taga (UC Berkeley), Dr. Jorge Escalante (University of Georgia), and Dr. Michael Thomas (UW-Madison) for thoughtful discussion on cobamide biosynthesis and advice on the development of microbiological detection assays, and members of the Kalan Laboratory for feedback.

## Author Contributions

Conceptualization, M.H.S and L.R.K.; Methodology, M.H.S, S.S., and L.R.K.; Formal Analysis, M.H.S., S.S.; Investigation, M.H.S, S.S., and L.R.K., Resources, L.R.K.; Data Curation, M.H.S., S.S., and L.R.K.; Writing – Original Draft, M.H.S., S.S., and L.R.K; Writing – Review & Editing, M.H.S. and L.R.K.; Visualization, M.H.S.; Funding Acquisition, M.H.S. and L.R.K.; Supervision, L.R.K.

## FIGURE LEGENDS

**Supplemental Figure 1. Read counts for cobamide biosynthesis genes, cobamide dependent genes, and rpoB**. The total sum of reads mapping to cobamide biosynthesis genes, cobamide dependent genes, or single-copy core gene rpoB within each sample are shown.

**Supplemental Figure 2. *De novo* cobamide biosynthesis is restricted to select bacterial families on the skin**. The top 20 abundant bacterial families within the dataset were determined by totaling the hits to single copy gene *rpoB* for each family. The remaining families were grouped into “Other”. Individual values in the heatmap represent the number of hits assigned to the family for a particular cobamide biosynthesis gene divided by the total number of hits to the gene. Gene hits were normalized by profile HMM coverage and sequencing depth prior to calculation. Black squares represent taxonomic abundance from 0 to 0.01%.

**Supplemental Figure 3.**. The number of unique species encoding single-copy core gene rpoB or 11 cobamide-dependent enzymes is shown. The number of unique *de novo* cobamide producers was determined by considering species with reads encoding for at least 5 of the 10 cobamide biosynthesis gene markers.

**Supplemental Figure 4.**. For each microenvironment network, the relative frequency of nodes with each given degree is shown.

**Supplemental Figure 5.**. Within each metagenome, the cumulative relative abundance of cobamide-producing Cutibacteria was calculated. A) NMDS plots based on Bray-Curtis indices for healthy adult samples within each skin microenvironment are shown. Points are colored by log 10 *Cutibacterium* cobamide producer abundance and sized by alpha diversity (Shannon).

**Supplemental Figure 6.**. Relative abundance of metagenomes with low or high *Corynebacterium* cobamide producer abundance. The first (0.05%) and third (0.75%) quartiles of the cobamide-producing Corynebacteria (CPC) relative abundance across all samples were used to group samples below 0.05% or above 0.75% CPC abundance. Relative abundances for samples within each group are shown. Species less than 10% relative abundance within each sample are grouped into “Other”.

**Supplemental Figure 7.**. Within each AD metagenome, the cumulative relative abundance of cobamide-producing Corynebacteria (CPC) was calculated. NMDS plots based on Bray-Curtis indices for pediatric AD samples within each skin microenvironment are shown. Points are colored by log 10 *Corynebacterium* cobamide producer relative abundance and sized by alpha diversity (Shannon). Shapes represent disease timepoint or healthy control.

**Supplemental Figure 8. Corynebacterium pangenome**. Pangenome analysis generated with anvi’o. 42,154 gene clusters (combined core, dispensable, and singletons) were identified from 71 Corynebacterium genomes and are ordered by gene cluster frequency (opaque, present; transparent, absent). Each gene cluster contains one or more genes contributed by one or more genomes. Genomes are colored by ecosystem association and ordered by the phylogeny based on 71 single copy genes (unrooted). ANI scale (0.7-0.8). Singleton gene clusters (grey) are collapsed.

**Supplementary Figure 9. *Corynebacterium* singleton gene clusters**. The number of singleton gene clusters per genome A) was determined using anvi’o. B) Welch’s unequal variances t-test did not reveal a significant difference in singleton gene cluster count between host- and environment-associated genomes.

**Supplemental Figure 10. *C. amycolatum* cell extract supports growth of *E. coli* strain auxotrophic for cobamides**. *E. coli* ATCC 14169 was used as a microbiological indicator for the detection of cobamide concentration in *C. amycolatum* LK19 cell extracts. A) Growth of *E. coli* was measured in minimal media with cyanocobalamin standards between 0.1 and 1.5 ng/mL to generate a standard curve. B) *E. coli* growth with cyanocobalamin standards or different dilutions of cell extract. OOD600 values from 6 biological replicates and at least 3 technical replicates are shown.

## METHODS

### Subject recruitment and sample collection

Healthy adult volunteers were recruited from the University of Wisconsin-Madison Microbial Sciences Building in Madison, WI, USA, from July through November 2019 under an institutional review board approved protocol. The single eligibility requirement was that the subject is over 18 years of age. Subjects provided written informed consent before participation. During each visit, 8 skin sites were sampled from that represent the physiologically diverse microenvironments of the skin: sebaceous (alar crease, occiput, back), moist (nare, antecubital fossa, umbilicus), dry (volar forearm), and foot (toe web space). Samples were collected by wetting a sterile foam swab (Puritan) with nuclease-free H_2_O and swabbing an approximately 1×1 inch area of the right lateral skin site for 15 rotations. Swabs were collected into 300 µL Lucigen MasterPure™ Yeast Cell Lysis solution and stored at -80°C until DNA extraction. Negative control air swabs, room swabs, extraction kit controls, and mock community samples were collected and prepared for sequencing as well.

For extraction, samples were thawed on ice and incubated shaking at 37°C for 1 hour in an enzymatic cocktail of ReadyLyse (Epicenter), mutanolysin (Sigma), and lysostaphin (Sigma). Swabs were then centrifuged in a filter tube insert (Promega) for 60 seconds at 21,300 x g to remove all liquid from the swab. The liquid was added to a glass bead tube (Qiagen) and vortexed for 10 minutes followed by incubation at 65°C shaking for 30 minutes and 5 minutes on ice. The liquid was removed and added to MPC protein precipitation reagent, vortexed thoroughly, and centrifuged for 10 minutes at 21,300 x g. The resulting supernatant was combined with isopropyl alcohol and column purified using the Invitrogen PureLink Genomic DNA extraction kit. Lastly, DNA was eluted in 50 µL of elution buffer.

### Metagenomic sequencing, processing, and taxonomic classification

Extracted DNA was prepared for sequencing by the University of Minnesota Genomics Center (UMGC) using the Nextera XT DNA Library Prep Kit (Illumina). Sequencing of the libraries was performed by UMGC on an Illumina NovaSeq (2 × 150 bp reads). We obtained 289 samples and 5.2 billion reads of non-human, quality-filtered, paired-end reads, with a median of 17.4 million paired-end reads per skin sample. Raw sequence data has been deposited in the NCBI Sequence Read Archive (SRA) under BioProject ID PRJNA763232.

Quality filtering, adapter removal, human decontamination, and tandem repeat removal were performed using fastp v0.21.0 (Chen et al., 2018) and KneadData v0.8.0. Taxonomic classification and abundance estimation was performed using Kraken 2 v2.0.8-beta (Wood et al., 2019) and Bracken v2.5 (Lu et al., 2017), with a custom database that included complete bacterial, viral, archaeal, fungal, protozoan, and human genomes, along with UniVec core sequences, from RefSeq. We further modified the custom database to separate plasmid sequences from the RefSeq genomes, as we and others have observed incorrect taxonomic assignment of plasmid sequences using RefSeq taxonomy with Kraken 2 (Doster et al., 2019). Potential contaminant species were identified and removed using the prevalence method in decontam v1.10.0 (Davis et al., 2018) and through manual inspection of air swab samples compared to matched skin swabs, ensuring a high-quality sequence set. To compare analyses across studies, an additional 906 human skin shotgun metagenomic samples from Oh *et al. (Oh et al*., *2016)* and Hannigan *et al. (Hannigan et al*., *2015)* were retrieved from the SRA under BioProject IDs PRJNA46333 and PRJNA266117, respectively. Metagenomic reads across multiple SRA run accessions from the same biological sample were pooled, processed for quality control, and assigned taxonomy using the methods outlined above. In all, the metagenomic data represents the skin microbial communities across 21 distinct sites from 66 healthy individuals. Sample information is described in Supplemental Material S1 and S2.

### Choice of profile HMMs for skin metagenome survey of cobamide biosynthesis and use

Profile HMMs, retrieved from the TIGRfam and Pfam databases, were used to detect cobamide biosynthesis, cobamide transport, and cobamide-dependent genes within skin metagenomic sequencing data. A total of 11 cobamide biosynthesis marker genes were selected because of their broad distribution throughout both the aerobic and anaerobic biosynthesis pathways and their presence within taxonomically-diverse cobamide producer genomes (Doxey et al., 2015; Lu et al., 2020; Shelton et al., 2019). CbiZ was included as a marker of cobamide remodeling, and btuB was included to assess cobamide transport. 19 cobamide-dependent enzymes and proteins with B_12_-binding domains were chosen to evaluate cobamide use. The single-copy core gene rpoB was used as a phylogenetic marker to assess microbial community structure within each metagenome and as a proxy for sequence depth. All cobamide-associated genes used in this analysis can be found in Supplemental Material S3.

### Metagenomic sequence search using HMMER

Sequencing reads from this study, Oh *et al*., and Hannigan *et al*. were converted to FASTA format, retaining only forward read files for analysis. For biological samples with multiple SRA run accessions, only the largest run when considering base pair count was included for analysis (Supplemental Material S1 and S2). Metagenomes were translated to each of 6 frame translations using transeq from the emboss v6.6.0 package (Rice et al., 2000). The program hmmsearch from HMMER v3.3.1 (Eddy, 2011) was used with default parameters and an E-value cutoff of 1E-06 to scan the metagenomic sequencing reads for homology to each cobamide-related HMM. The resulting hits were taxonomically classified to the species level using Kraken 2 (Wood et al., 2019) and Bracken (Lu et al., 2017). The number of hits for each gene was normalized to HMM length when analyzing individual metagenomes and to both HMM length and sequencing depth when analyzing groups of metagenomes. To reduce the rate of rare and singleton hits, species-gene pairings that did not appear in at least five samples from two or more datasets were excluded from further analysis. Taxonomic frequency profiles were generated for each cobamide-related gene by dividing the normalized number of gene hits per taxon by the total normalized number of gene hits.

### Metagenomic sequence search using INFERNAL

Covariance models (CMs) for 3 cobalamin riboswitches from the Rfam clan CL00101 were retrieved from the Rfam database (Kalvari et al., 2018) (Supplemental Material S3). The program cmsearch from INFERNAL v1.1.2 (Nawrocki and Eddy, 2013) was used with default parameters and an E-value cutoff of 1E-06 to scan the metagenomes for RNA homologs to cobalamin riboswitches. The methods following hit identification are the same as described above for HMM analysis, except that the number of hits for each riboswitch were not normalized by CM length because the read lengths and CM lengths were relatively similar.

### Mapping of cobalamin riboswitch hits to genomes

Reads identified from INFERNAL were aligned against the complete genomes of *Cutibacterium acnes* KPA171202, *Veillonella parvula* DSM 2008, *Pseudomonas putida* KT2440, *Corynebacterium amycolatum* FDAAROS 1107, and *Streptococcus sanguinis* SK36 using bowtie2 v2.3.5.1 (Langmead and Salzberg, 2012) and visualized in R with the ggbio v1.30.0 package (Yin et al., 2012). Genes upstream and downstream of the aligned reads within each genome were assigned functions based on NCBI RefSeq annotations and visualized using the gggenes v0.4.0 R package. Genes within genomic regions that encoded for a cobalamin riboswitch but had no genes currently known to be under cobalamin riboswitch control were assigned putative functions based on the top hit from NCBI BLAST searches against the nt/nr nucleotide collection database.

### SPIEC-EASI microbial network inference

The statistical method SPIEC-EASI from the SpiecEasi R package v1.0.7 (Kurtz et al., 2015) was used to identify associations between microbial species in the skin metagenomes. Samples included for analysis are indicated in Supplemental Material S1 and S2; samples with comparatively low read counts within each dataset were excluded. Species were included for analysis if they were present at greater than 0.015% average abundance and identified in at least 55% of the samples, resulting in 185 final species Supplemental Material S5). SPIEC-EASI (neighborhood selection mode) was performed on samples grouped by both microenvironment and study, resulting in 12 total networks. Consensus networks were created for each microenvironment by merging sign-consistent edges from the node and edge sets identified for each study, requiring that each edge appear in at least 2 of the 3 datasets (Kurtz et al., 2019). The R package igraph v1.2.6 (Csardi et al., 2006) was used for network visualization and calculation of topological network properties.

To incorporate cobamide biosynthesis and dependence information into the network analysis, data from Supplementary Table 5 of Shelton *et al. (Shelton et al*., *2019)* was used to assess the presence of cobamide dependent enzymes and the potential for cobamide biosynthesis or precursor salvage in each species in the consensus networks. For each species, the presence of 7 cobamide-dependent enzymes (‘B12-dependent RNR’, ‘metH’, ‘methylmalonyl-CoA mutase family’, ‘ethanolamine ammonia lyase’, ‘B12-dependent glycerol/diol dehydratase’, ‘D-ornithine 4,5-aminomutase’, ‘epoxyqueosine reductase’) was determined, and the ‘cobamide biosynthesis category’ was used to assign cobamide biosynthesis potential. These 7 cobamide dependent enzymes were chosen because they represent those most abundant on the skin (Figure 3). For any consensus network species absent from the Shelton *et al*. dataset, KO identifiers in Supplemental Material S6 were searched against NCBI RefSeq complete assembled genomes for each species using KOfamScan, a functional annotation program based on KOs and HMMs (Aramaki et al., 2020). Genomes were then scored for cobamide biosynthesis category based on the presence of certain sets of cobamide biosynthesis genes (Supplemental Material S5 and S6).

### Microbiome diversity analysis of *Corynebacterium* cobamide producers

For microbiome diversity analyses of the healthy adult skin microbiome, metagenomes from this study and Oh *et al*. (Oh et al., 2016) were subsampled to 1.5 million read counts using rarefy_even_depth() from the phyloseq R package v1.34.0 (McMurdie and Holmes, 2013), discarding samples below this read count cutoff. Samples included for analysis are indicated in Supplemental Material S1 and S2; samples from Hannigan *et al*. (Hannigan et al., 2015) were excluded from analysis due to comparatively lower sequencing depth; median 1.2 million (Hannigan) vs 17.4 million (this study) and 16.9 million (Oh) final paired-end reads. To adjust for study effect, adjust_batch() from the MMUPHin R package v1.5.2 (Ma, 2021) was used. Using taxonomic abundance information, the cumulative relative abundance of *Corynebacterium* species that encode for *de novo* cobamide biosynthesis was calculated for each metagenome. Alpha diversity was determined by calculating the Shannon index using the phyloseq diversity() function. For beta diversity analysis, abundances were square root transformed to give more weight to low abundance taxa, and the Bray-Curtis dissimilarity index was calculated for samples within each skin site using vegdist() from the vegan v2.5-6 package. The indices were ordinated using non-metric multidimensional scaling with the vegan metaMDS() program. For analysis of the pediatric atopic dermatitis microbiome, metagenomes from Byrd *et al*. (Byrd et al., 2017) were accessed from the SRA, processed, and assigned taxonomy using the described methods (Supplemental Material S7). Samples MET1440, MET1441, MET1449, MET1552, and MET1563 were excluded due to insufficient sequence data after processing. Analysis of *Corynebacterium* cobamide producer abundance and alpha and beta diversity was performed as outlined above.

### *Corynebacterium* comparative genomics

71 *Corynebacterium* isolate genomes were acquired either from the National Center for Biotechnology Information (NCBI) as complete assemblies or from human skin isolates as draft assemblies. Supplemental Material S8 reports accession numbers and other information for each isolate genome, including ecosystem association, which was assigned using strain metadata and species-specific literature. The pangenomics workflow from anvi’o v6.2 (http://merenlab.org/2016/11/08/pangenomics-v2/) (Delmont and Murat Eren, 2018; Eren et al., 2015) was used for comparative genomics analysis. Briefly, genomes were annotated using ‘anvi-run-ncbi-cogs’, which assigns functions from the NCBI Clusters of Orthologous Groups (COGs) database. The *Corynebacterium* pangenome was computed using the program ‘anvi-pan-genome’ with the flags ‘--minbit 0.5’, --mcl-inflation 6’, and ‘--enforce-hierarchical-clustering’. Average nucleotide identity between genomes was calculated using pyani within the anvi’o environment (https://github.com/widdowquinn/pyani) (Pritchard et al., 2015). The program ‘anvi-get-enriched-functions-per-pan-group’ was utilized to identify enriched COGs between host- and environment-associated genomes (Shaiber et al., 2020). Genome summary statistics are presented in Supplemental Material S8.

### *Corynebacterium* phylogenetic analysis

The anvi’o phylogenomics workflow (http://merenlab.org/2017/06/07/phylogenomics/) was used to create a *Corynebacterium* phylogeny. Within the anvi’o environment, single-copy core genes (SCGs) from the curated anvi’o collection Bacteria_71 were identified within each genome using HMMER (Eddy, 2011), and the SCG amino acid sequences were concatenated and aligned using MUSCLE (Edgar, 2004). A phylogenetic tree was then constructed using FastTree (Price et al., 2010) within the anvi’o environment, and *Tsukamurella paurometabola* DSM 20162 was included as an outgroup to root the tree. To identify cobamide biosynthesis genes within the 71 *Corynebacterium* genomes, KEGG orthology (KO) identifiers from KEGG map00860 (Supplemental Material S6) were used to create a custom profile for KOfamscan. Each genome was queried against this profile, and hits to the KOs above the predefined inclusion threshold or user-defined threshold (cobU/cobT, cobC, and bluB), were considered for further analysis. Visualization of the phylogenetic tree and cobamide biosynthesis pathway completeness was performed in R with the ggtree package v2.4.2 (Yu et al., 2017).

### Corynebacterium minimal M9 (CM9) medium composition and preparation

11.28 g/L M9 salts, 0.1 g/L L-dextrose, and 0.2% Tween 80 were prepared in aqueous solution and autoclaved at 121°C for 15 minutes. We and others have observed that autoclaving a small amount of L-dextrose in the presence of other media components improves rapid and abundant growth of *Corynebacterium* species in synthetic media (Liebl et al., 1989). When cooled, the following media components were added at the concentrations indicated: 0.1 mM CaCl_2_, 2 mM MgSO_4_, 50 nM CoCl_2_, 6 µM thiamine-HCl, 1.9 g/L L-dextrose, 100 mg/L L-arginine, 2 mg/L biotin.

### *Corynebacterium amycolatum* cell extract preparation

*C. amycolatum* LK19, isolated from healthy adult skin, was cultured overnight in BHI with 0.2% Tween 80. To remove residual cobamides in the media and scale up culture conditions, cells were washed 3 times with CM9 broth, inoculated into 250-mL CM9 broth (starting OD600 = 0.1), and incubated shaking at 37°C for 24 hours. Cells were again washed, inoculated into 1-L CM9 broth (starting OD600 = 0.1), and incubated shaking at 37°C for 48 hours. Cells were spun down at 4000 rpm for 20 minutes, wet cell weight was recorded, and 20 mL methanol per 1 g wet cell weight was added for metabolite extraction. To convert cobamides to their cyano form, 20 mg potassium cyanide was added per 1 g wet cell weight, and the cell suspension was heated at 60°C for 90 minutes and mixed intermittently every 20 minutes. Following overnight room temperature incubation, cell debris was removed, and the solvent was evaporated using a rotary evaporator. The resulting extract was de-salted with a C18 Sep-Pak (Waters) cartridge. Briefly, the cell extract was suspended in 10-ml H_2_O and run through the cartridge, followed by a 20-mL H_2_O wash and elution of the cobamide-containing fraction with 3-mL methanol. The desalted extract was dried in the fume hood and resuspended in 1.1 mL H_2_O for subsequent analysis.

### Microbiological cobamide indicator assay

*E. coli* strain ATCC 14169, which requires either cobamide or methionine supplementation for growth, was acquired from the NRRL Culture Collection. The strain was cultured for 6 hours in BHI with 0.2% Tween 80 and washed 3 times with M9 minimal medium (11.28 g/L M9 salts (Sigma-Aldrich), 0.1 mM CaCl_2_, 0.2 mM MgSO_4_, 1 mM thiamine-HCl, 2 g/L L-dextrose, 100 mg/L L-arginine). Cells were adjusted to an OOD600 of approximately 0.02 in M9 minimal medium. In each well of a 96-well plate, 200 uL of cells and 2.5 uL of sample (cyanocobalamin standards or *C. amycolatum* LK19 cell extract dilutions) were added. The plate was incubated stationary at 37°C for 18 hours, and OOD600 values were recorded using a BioTek Epoch 2 Microplate Spectrophotometer. A standard curve was generated using cyanocobalamin concentrations between 0.1 and 1.5 ng/mL, and this was used to calculate cobamide concentration in the cell extracts. Intracellular concentrations were estimated assuming a cellular volume of 1 µm^3^ and 8×10^8^ cells/mL at an OD_600_ of 1.0 (Sokolovskaya et al., 2019).

### Quantification and Statistical Analysis

The R Statistical Package was used to generate figures and compute statistical analyses. Statistical significance was verified through the non-parametric Wilcoxon rank-sum test with FDR correction or Welch’s unequal variances t-test. Correlations between cobamide-producing Corynebacteria abundance and Shannon diversity were calculated using the Spearman rank coefficient.

