## Supplemental Figures 1-10 for "Cobamide sharing drives skin microbiome dynamics"

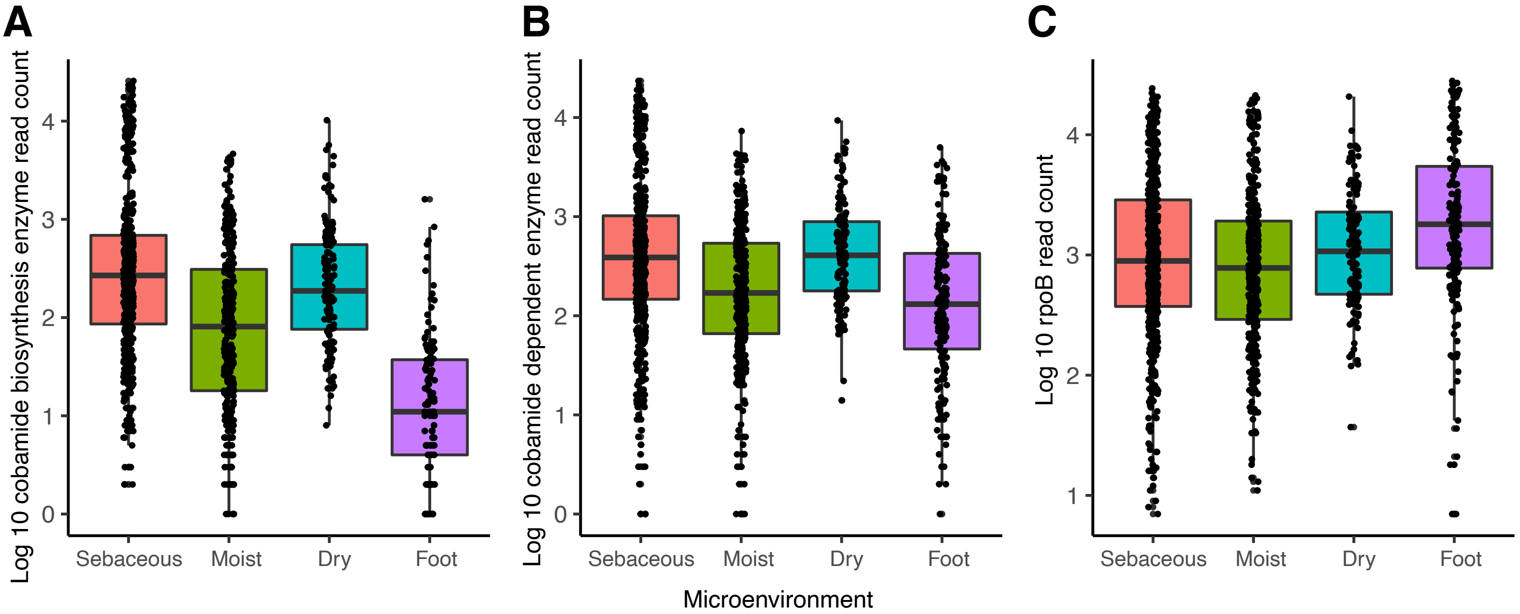


**Supplemental Figure 1. Read counts for cobamide biosynthesis genes, cobamide dependent genes, and rpoB.** The total sum of reads mapping to cobamide biosynthesis genes, cobamide dependent genes, or single-copy core gene rpoB within each sample are shown.

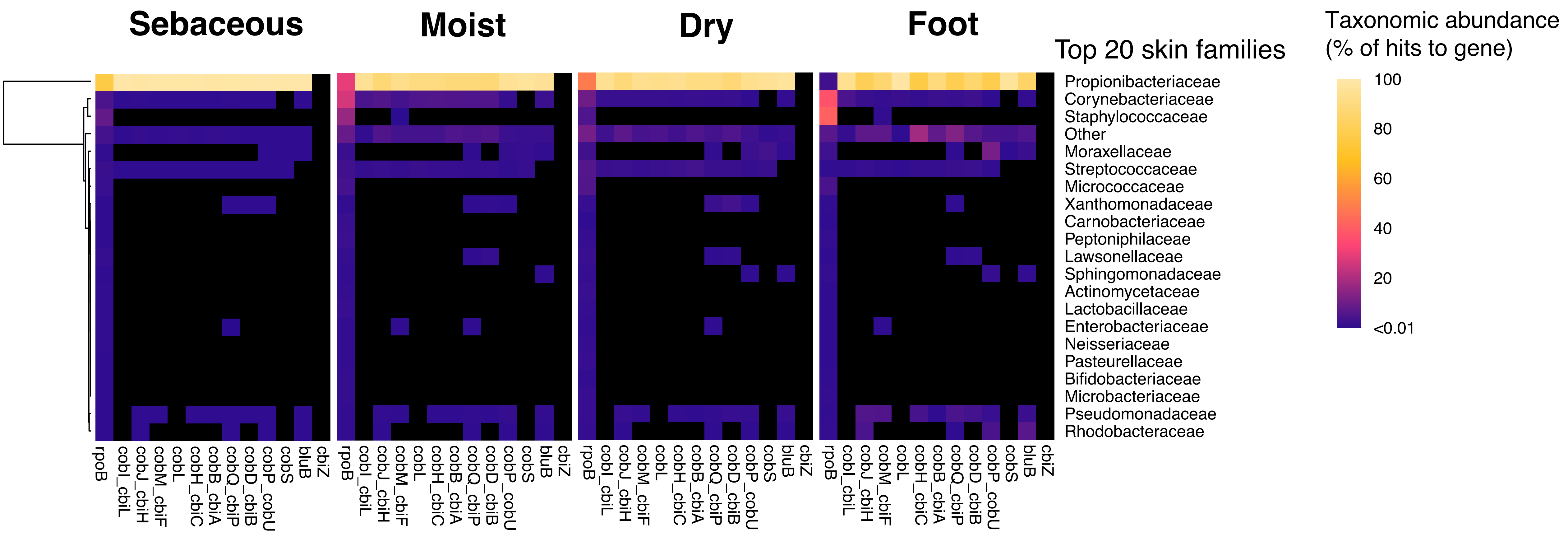


**Supplemental Figure 2. De novo cobamide biosynthesis is restricted to select bacterial families on the skin.** The top 20 abundant bacterial families within the dataset were determined by totaling the hits to single copy gene rpoB for each family. The remaining families were grouped into “Other”. Individual values in the heatmap represent the number of hits assigned to the family for a particular cobamide biosynthesis gene divided by the total number of hits to the gene. Gene hits were normalized by profile HMM coverage and sequencing depth prior to calculation. Black squares represent taxonomic abundance from 0 to 0.01%.

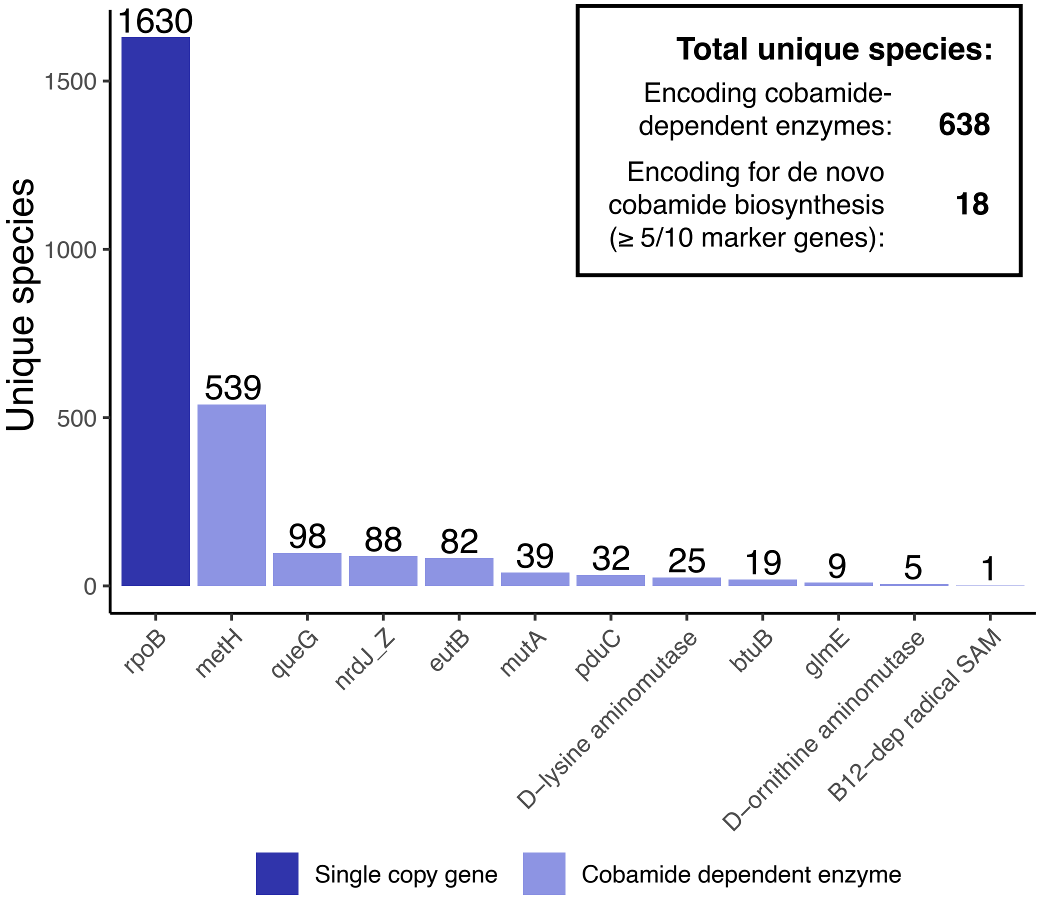


**Supplemental Figure 3.** The number of unique species encoding single-copy core gene rpoB or 11 cobamide-dependent enzymes is shown. The number of unique de novo cobamide producers was determined by considering species with reads encoding for at least 5 of the 10 cobamide biosynthesis gene markers.

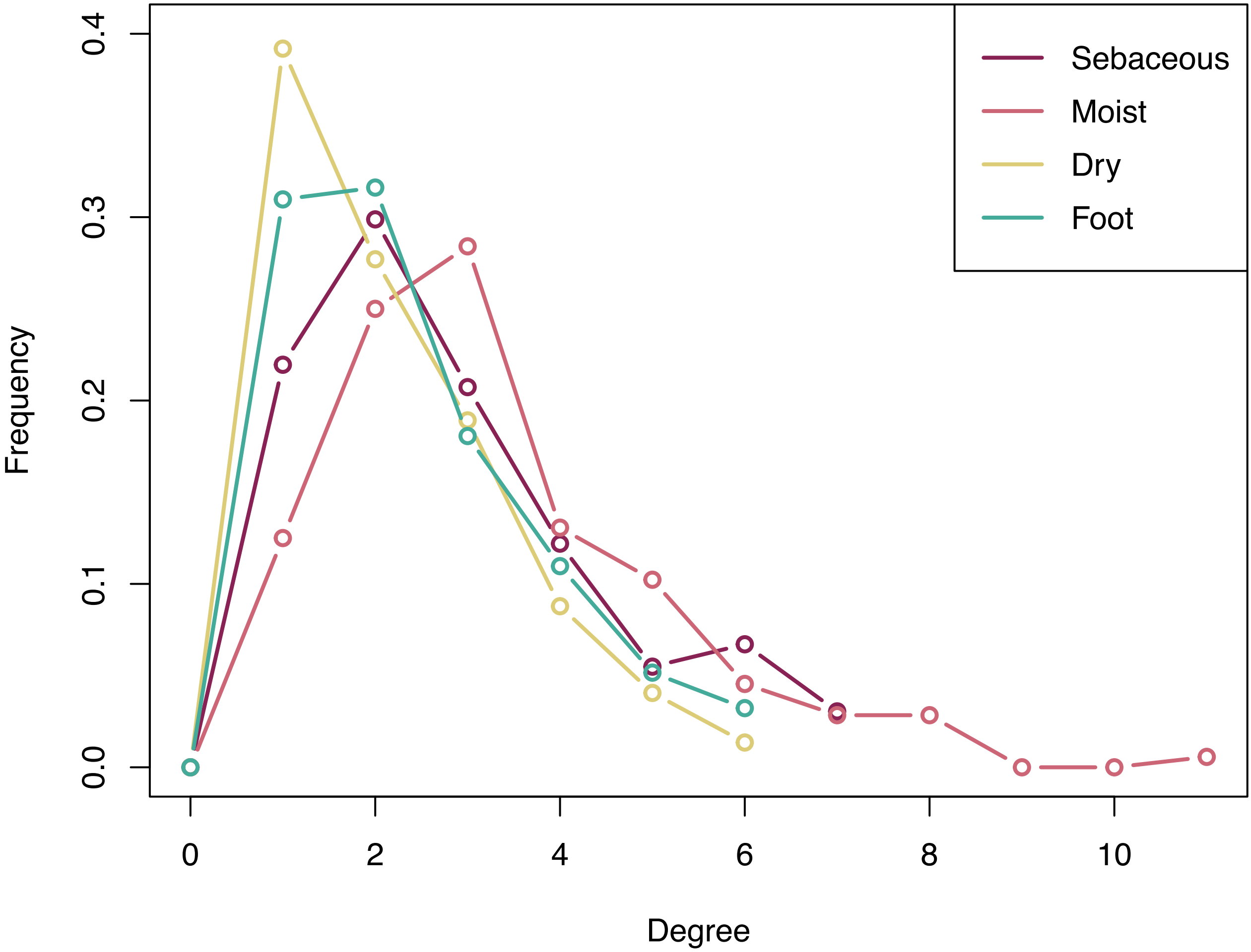


**Supplemental Figure 4.** For each microenvironment network, the relative frequency of nodes with each given degree is shown.

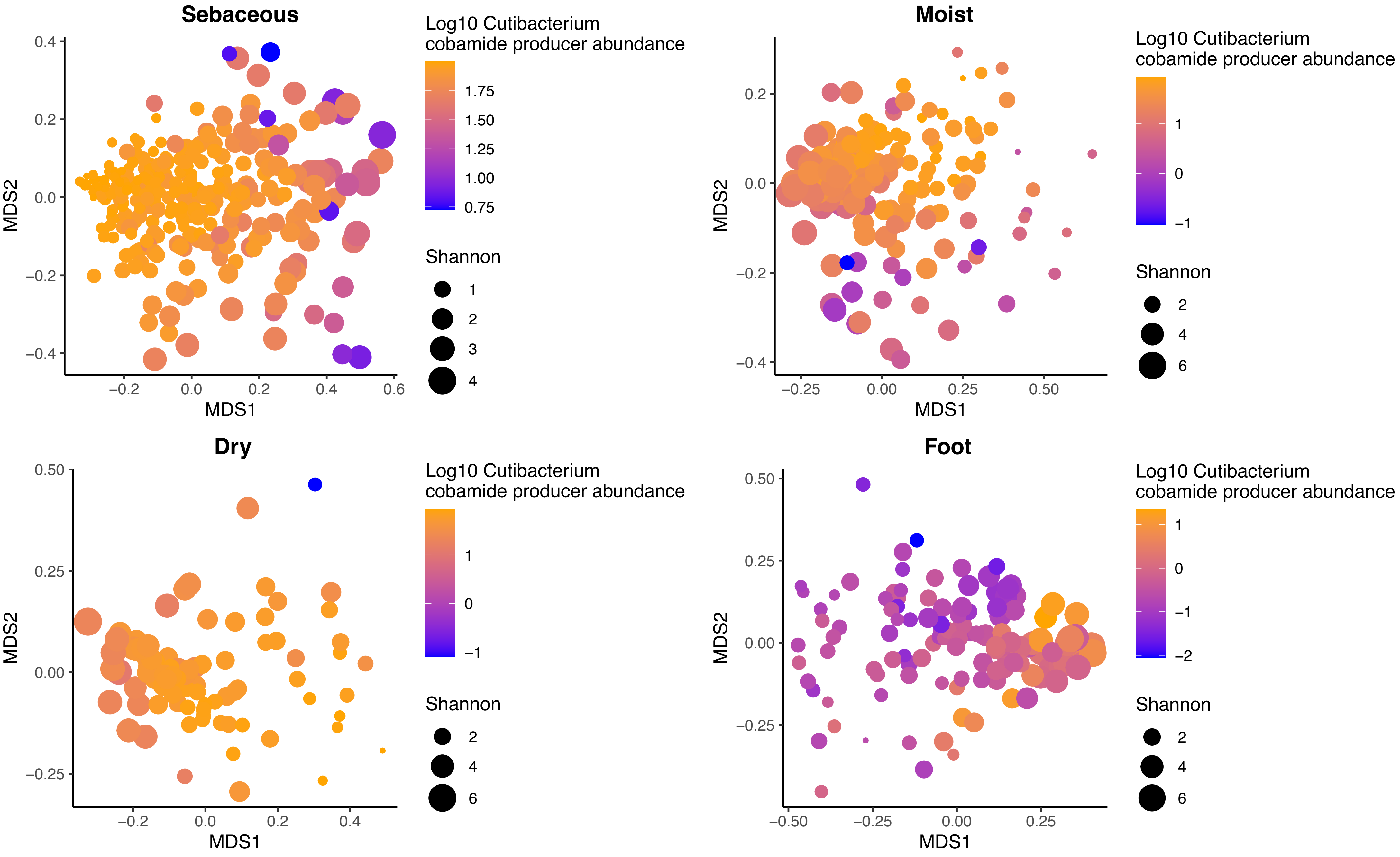


**Supplemental Figure 5.** Within each metagenome, the cumulative relative abundance of cobamide-producing Cutibacteria was calculated. A) NMDS plots based on Bray-Curtis indices for healthy adult samples within each skin microenvironment are shown. Points are colored by log 10 Cutibacterium cobamide producer abundance and sized by alpha diversity (Shannon).

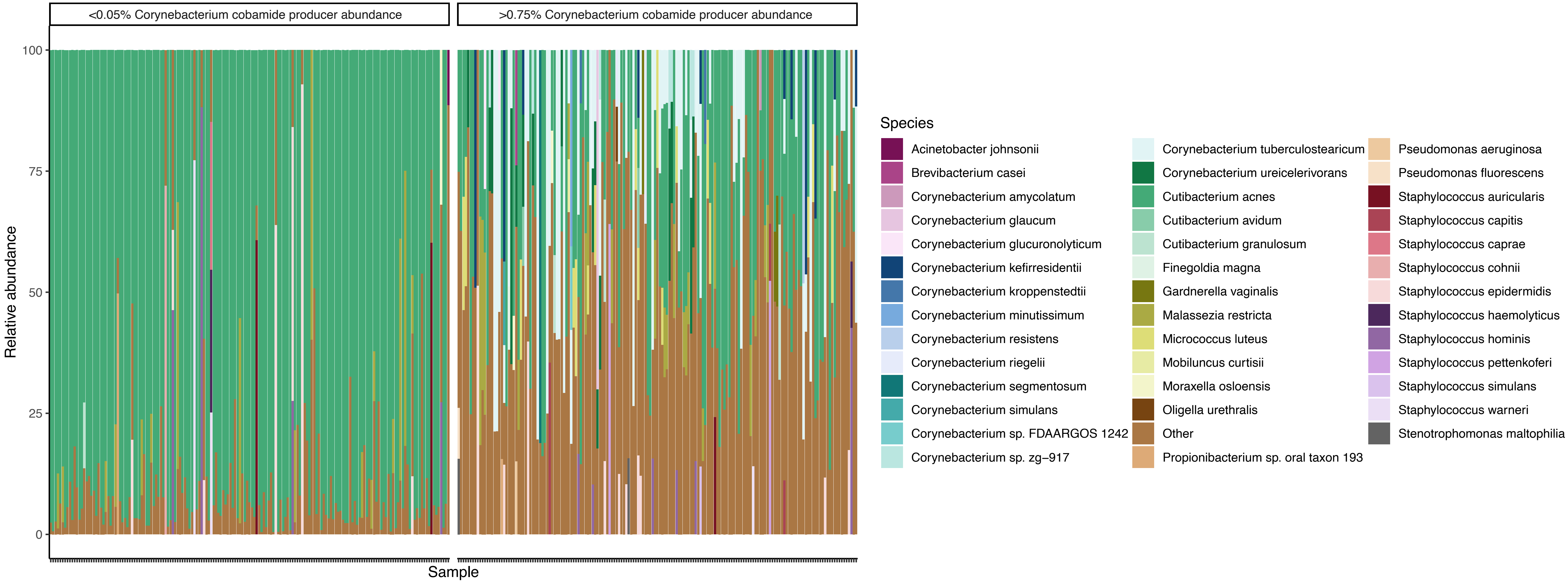


**Supplemental Figure 6. Relative abundance of metagenomes with low or high Corynebacterium cobamide producer abundance.** The first (0.05%) and third (0.75%) quartiles of the cobamide-producing Corynebacteria (CPC) relative abundance across all samples were used to group samples below 0.05% or above 0.75% CPC abundance. Relative abundances for samples within each group are shown. Species less than 10% relative abundance within each sample are grouped into “Other”.

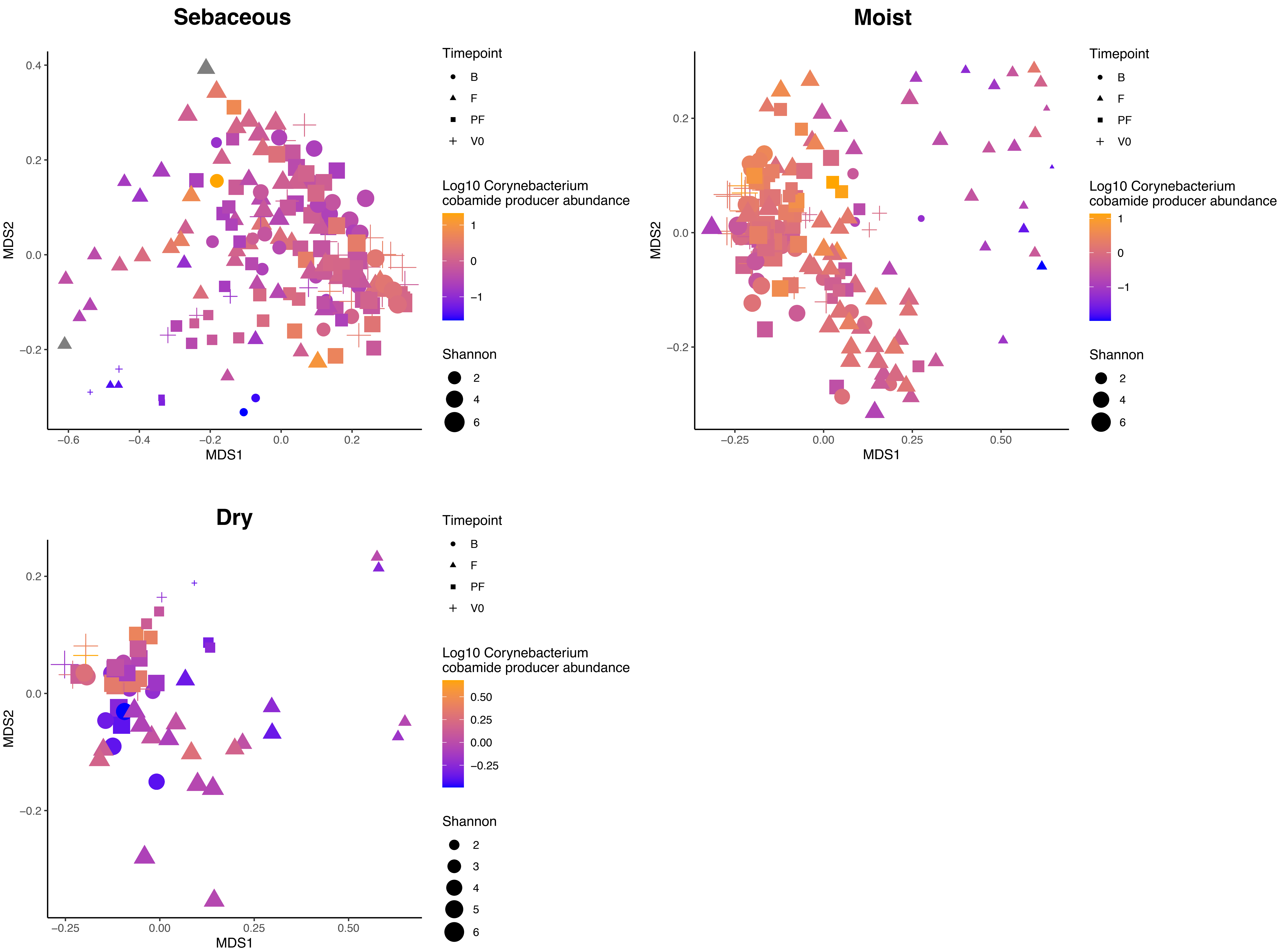


**Supplemental Figure 7.** Within each AD metagenome, the cumulative relative abundance of cobamide-producing Corynebacteria (CPC) was calculated. NMDS plots based on Bray-Curtis indices for pediatric AD samples within each skin microenvironment are shown. Points are colored by log 10 Corynebacterium cobamide producer relative abundance and sized by alpha diversity (Shannon). Shapes represent disease timepoint or healthy control.

### Corynebacterium Pangenome

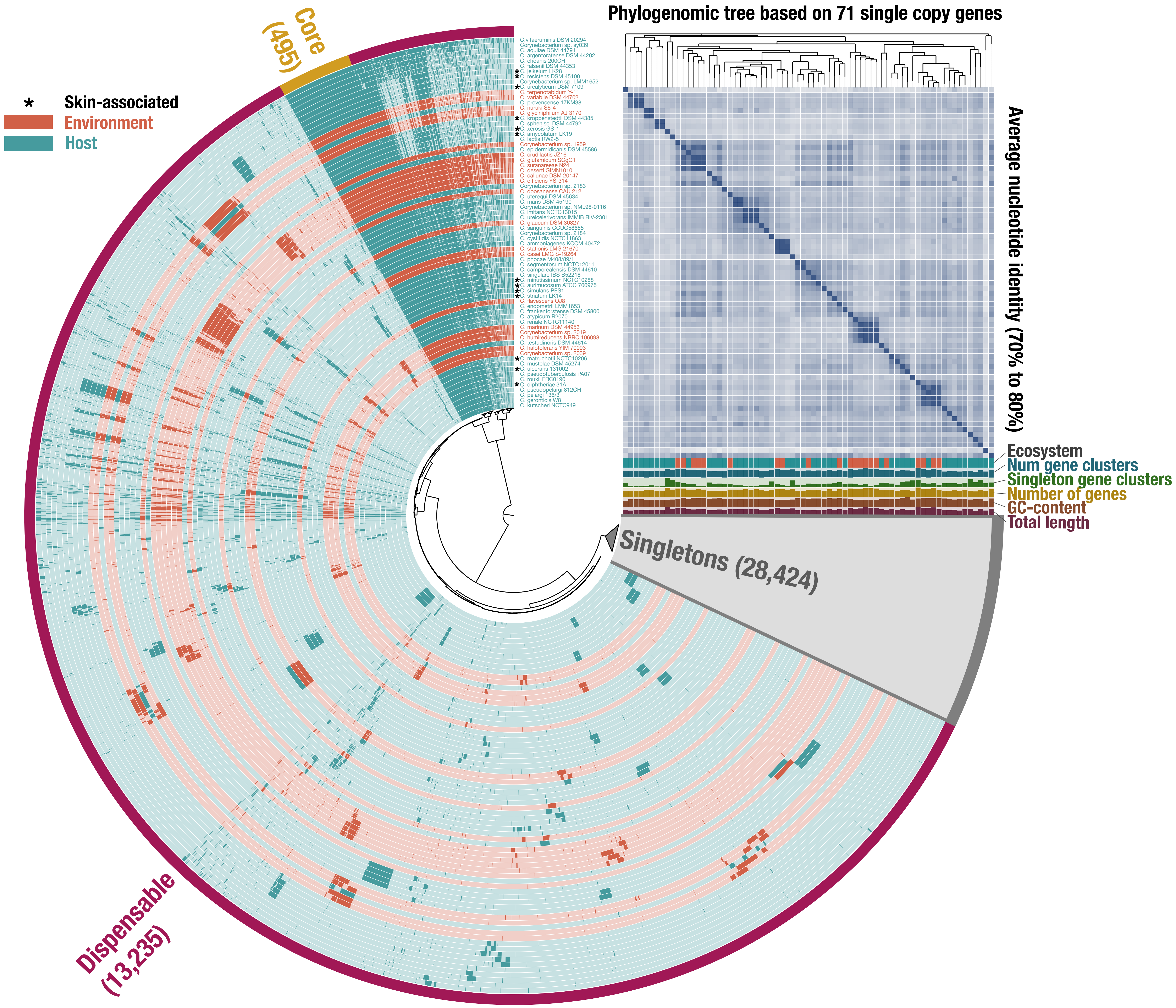


**Supplemental Figure 8. Corynebacterium pangenome.** Pangenome analysis generated with anvi'o. 42,154 gene clusters (combined core, dispensable, and singletons) were identified from 71 Corynebacterium genomes and are ordered by gene cluster frequency (opaque, present; transparent, absent). Each gene cluster contains one or more genes contributed by one or more genomes. Genomes are colored by ecosystem association and ordered by the phylogeny based on 71 single copy genes (unrooted). ANI scale (0.7-0.8). Singleton gene clusters (grey) are collapsed.

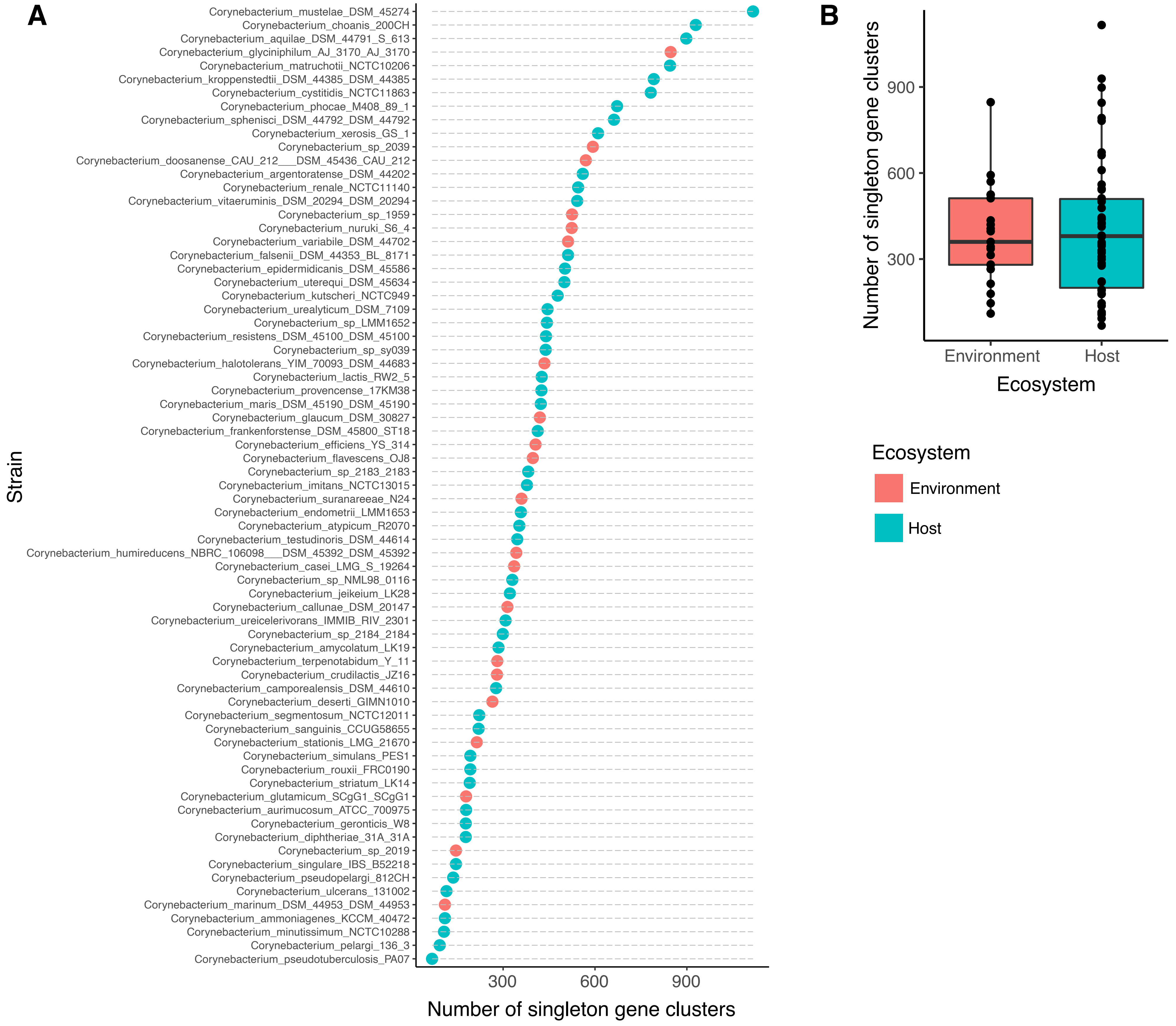


**Supplementary Figure 9. Corynebacterium singleton gene clusters.** The number of singleton gene clusters per genome A) was determined using anvi'o. B) Welch's unequal variances t-test did not reveal a significant difference in singleton gene cluster count between host- and environment-associated genomes.

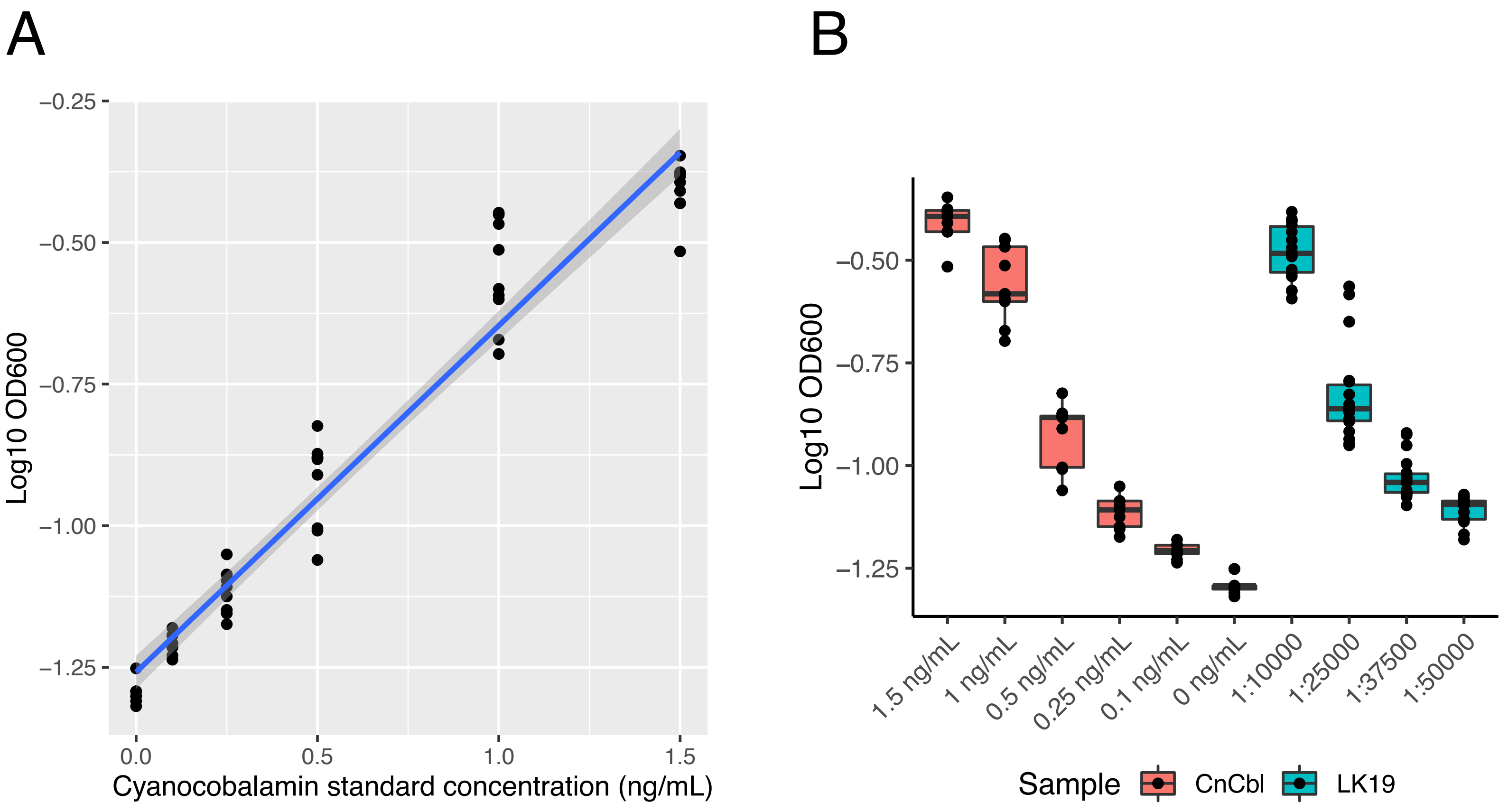


**Supplemental Figure 10. C. amycolatum cell extract supports growth of E. coli strain auxotrophic for cobamides.** E. coli ATCC 14169 was used as a microbiological indicator for the detection of cobamide concentration in C. amycolatum LK19 cell extracts. A) Growth of E. coli was measured in minimal media with cyanocobalamin standards between 0.1 and 1.5 ng/mL to generate a standard curve. B) E. coli growth with cyanocobalamin standards or different dilutions of cell extract. OOD600 values from 6 biological replicates and at least 3 technical replicates are shown.
