## Supplemental Tables 1-2 for "Cobamide sharing drives skin microbiome dynamics"

### Supplemental Table 1. Properties of skin microenvironment networks.

| Microenvironment | Edges | Nodes | Density* | Transitivity** | Phylum Assortativity*** | Modularity**** | Average node degree ***** |
|---|---|---|---|---|---|---|---|
| **Sebaceous** | 231 | 164 | 0.017283 | 0.2187005 | 0.7036747 | 0.7598433 | 2.82 |
| **Moist** | 288 | 176 | 0.018701 | 0.2381974 | 0.7059118 | 0.7444059 | 3.27 |
| **Dry** | 159 | 148 | 0.014617 | 0.1843003 | 0.6873054 | 0.8310193 | 2.15 |
| **Foot** | 184 | 155 | 0.015417 | 0.2 | 0.8162985 | 0.8135633 | 2.37 |

*Density represents the number of existing relationships relative to the total possible number
** Transitivity indicates the probability that two adjacent nodes are connected and can reveal the existence of tightly connected communities
*** Phylum assortativity reflects the preference for nodes to be connected to other nodes within the same phylum
**** Modularity is a measurement of the division of a network into modules
***** Average node degree is a measure of network sparsity; a lower average node degree indicates a sparser network.

**Supplemental Table 2. Spearman correlation analysis of Log 10 cobamide-producing *Corynebacterium* abundance and Shannon diversity.**

| Microenvironment | Rho value | P-value |
|---|---|---|
| **Sebaceous** | 0.63 | < 2.2e-16 |
| **Moist** | 0.46 | 1.07e-10 |
| **Dry** | 0.14 | 0.20 |
| **Foot** | 0.21 | 0.02 |
